# Functional characterization of a cytosolic malic enzyme crucial for pyridine nucleotide homeostasis, redox balance, and virulence in *Leishmania major*

**DOI:** 10.64898/2026.07.14.735955

**Authors:** Arpita Pal, Debasmita Modak, Faraz Khan, Dipon Kumar Mondal, Samudrala Gourinath, Rupak Datta

**Affiliations:** Department of Biological Sciences, Indian Institute of Science Education and Research Kolkata, Mohanpur, West Bengal, India; Structural Biology Laboratory, School of Life Sciences, Jawaharlal Nehru University, New Delhi, India

**Keywords:** Leishmania major, malic enzyme, pyridine nucleotide homeostasis, reactive oxygen species, virulence

## Abstract

Malic enzymes (MEs) occupy a central node in metabolism, by catalyzing the reversible oxidative decarboxylation of malate to pyruvate, thereby contributing to maintenance of NADPH homeostasis. *Leishmania spp.* are known to encode two ME isoforms, however only one from *Leishmania major* has been functionally characterised till date to be a mitochondrial enzyme, playing critical role in gluconeogenesis. Here, we cloned and functionally characterized the second isoform, LmME2. Colocalization and subcellular fractionation analysis established that LmME2 localizes to the cytosol. Kinetic analyses of purified LmME2 revealed comparable rates of malate decarboxylation and pyruvate carboxylation, however, pyruvate carboxylation predominated in parasite cell lysates, indicating that the directionality of the reaction is metabolically regulated in cellular environment. Consistent with this, we identified oxaloacetate, ATP and fumarate as allosteric modulators of LmME2 activity. To dissect its physiological role, we generated CRISPR-Cas9-mediated LmME2 knockout parasites (LmME2^-/-^). Interestingly, LmME2 deletion did not alter NADPH levels but depleted both NADP^+^ and NAD^+^ pools, with reduction in NAD^+^ being more pronounced. This disruption of pyridine nucleotide homeostasis resulted in elevated intracellular ROS levels. Furthermore, inhibition of the pentose phosphate pathway, the alternative cytosolic source of NADPH, severely impaired the growth of LmME2^-/-^ strain, highlighting the role of LmME2 in maintaining parasite fitness. These metabolic defects translated into markedly diminished intracellular survival of LmME2^-/-^ parasites and its attenuated virulence in mice. Collectively, our findings identify LmME2 as the first functionally characterized cytosolic malic enzyme in *Leishmania* and establish it as a redox-associated virulence factor. These results further highlight LmME2 as a promising antileishmanial drug target.

## Introduction

*Leishmania spp.* are kinetoplastid protozoan parasites of the family Trypanosomatidae that are transmitted by phlebotomine sandflies and cause leishmaniasis. The disease manifests as cutaneous, mucocutaneous, or visceral forms depending on the infecting species, collectively affecting millions of people worldwide (1). Limitations of currently available treatments, including toxicity, prolonged treatment regimens, high cost, and emerging drug resistance, call for a better understanding of parasite biology to guide drug development (2, 3). The parasite has a digenetic life cycle: during a blood meal, an infected sandfly transfers motile promastigotes to the mammalian host, where they transform into amastigotes after being taken up by macrophages (4, 5). The two life forms thus inhabit markedly different environments with distinct challenges. In the sandfly gut, promastigotes reside in a nutrient-replete milieu but are exposed to oxidative stress arising from digestion of heme-rich blood meal (6). In contrast, amastigotes survive and proliferate within the mammalian macrophage phagolysosome (7–9), where they encounter a hostile environment characterized by acidic pH, oxidative burst, and restricted availability of sugars and essential micronutrients (10–14). Adaptation to these fluctuating conditions depends on a specialised metabolic network, and the metabolic players that underpin it are therefore of interest both as determinants of parasite survival and as candidate drug targets.

Malic enzymes (MEs) are oxidoreductases that catalyse the reversible oxidative decarboxylation of malate to pyruvate, coupled to the reduction of NAD(P)+ to NAD(P)H (15). First isolated from pigeon liver by Ochoa, Mehler, and Kornberg in 1947 (16), MEs have since been identified across all domains of life. Although the reaction is reversible, physiological functions are predominantly associated with malate decarboxylation (17–19). However, there are a few contexts in which pyruvate carboxylation was found to be operative (20, 21). Mammals possess three ME isoforms. The cytosolic ME1 primarily supports fatty acid biosynthesis in lipogenic tissues and maintains cellular redox homeostasis through NADPH generation via malate decarboxylation (22, 23). ME1 has also been reported to operate in the reverse pyruvate carboxylating direction in hypertrophied rat heart, where it provides anaplerotic input to the TCA cycle (24). Of the two mitochondrial isoforms, ME2 contributes to mitochondrial malate-pyruvate cycling through the generation of pyruvate and reducing equivalents that support mitochondrial respiration (25, 26), while ME3 has been shown to be important for glucose-stimulated insulin secretion in pancreatic β-cells through a mechanism which may not depend primarily on its catalytic activity (27). The physiological importance of MEs has been particularly well characterised in cancer cells. ME1 and ME2 are frequently upregulated in tumours, where they support proliferation by supplying NADPH for reductive biosynthesis and redox balance and by channelling carbon between glycolysis and the TCA cycle (17, 28, 29). These findings have established MEs as important regulators of metabolic adaptation and have attracted interest in their therapeutic targeting (26).

In plants, distinct ME isoforms have been identified in cytosol, chloroplasts and mitochondria (30, 31). Through malate decarboxylation, chloroplastic MEs supply COc for photosynthesis in C4 species (32), whereas mitochondrial MEs were reorted to have a major impact on nocturnal metabolism (31). Bacterial genomes typically encode between one and four malic enzymes - for example, *Escherichia coli* carries two paralogues (MaeA and MaeB) (33), while *Bacillus subtilis* possesses four enzymes of differing cofactor preference (34). Among fungi, the single mitochondrial ME of *Saccharomyces cerevisiae* (35) contrasts with the multiple isoforms found in oleaginous species (19), which have been exploited for biotechnological lipid production through increased NADPH availability (36).

Despite these advances, the functions of MEs in trypanosomatid parasites remain poorly understood. Biochemical studies have established that the MEs of *Trypanosoma cruzi* and *Trypanosoma brucei* are NADP^+^-dependent enzymes with distinct cytosolic and mitochondrial localisations (37). In terms of physiological role, the cytosolic ME in *T. brucei* was reported to act as a redundant route to pentose phosphate pathway in countering oxidative stress during glucose limitation (38). In contrast, the ME orthologs in *Leishmania* remain largely unexplored. Comparative genomic analyses suggested that most *Leishmania* species encode two ME isoforms (39). Although biochemical and kinetic analyses have been reported previously for two ME homologues, one each from *L. major* and *L. mexicana* (39), the study did not report their functional characterisation. To date, only a mitochondrial ME (hereafter referred to as LmME1) has been functionally characterised by us recently in *L. major*, where it was shown to play an important role in parasite gluconeogenesis through its pyruvate carboxylating activity (21). However, the second ME isoform remains completely uninvestigated; neither its localisation or its physiological role has been established in any *Leishmania* species. In the present study, we characterize the second malic enzyme isoform in *L. major*, designated LmME2. We demonstrate that LmME2 is a catalytically active NADP^+^-dependent malic enzyme that localizes to the cytosol and is subject to metabolite-mediated regulation. Genetic deletion of LmME2 depleted NADP^+^ and NAD^+^ levels, elevated intracellular ROS levels, and compromised parasite survival within macrophages as well as in a murine infection model. Together, our findings provide the first functional characterisation of a cytosolic malic enzyme in *Leishmania* and establish LmME2 as a determinant of parasite virulence through maintenance of redox balance by contributing to pyridine nucleotide homeostasis.

## Results

### LmME2 is expressed in both promastigote and amastigote stages of *Leishmania major*

*Leishmania* species are predicted to encode two malic enzyme isoforms. In the revised *L. major* genome assembly (OU755558.1) (40), two ME isoforms are annotated on Chromosome 24 as LMJFC_240013800 and LMJFC_240013900, which are located in tandem at positions 268814-270529 bp and 274438-276159 bp respectively (Fig. 1A). Notably, LMJFC_240013800 is flanked by repetitive SIDER elements, suggesting that the two genes may have arisen through a duplication event (Fig. 1A). Our previous work demonstrated that LMJFC_240013900 (LmME1) encodes a mitochondrial malic enzyme that supports gluconeogenesis in *L. major* (21). However, the expression, localisation, and physiological role of the second paralogue, LMJFC_240013800 (referred to as LmME2), have not been experimentally investigated. To validate the presence of the LmME2 locus in *L. major* genome, full-length primers (P1, P2) were designed against the coding region of LMJFC_240013800. PCR amplification with these primers with the *L. major* genomic DNA yielded the expected ∼1.7 kb product (Fig. 1B), suggesting the presence of an ME2 isoform in the *L. major* genome. Sequencing of the LmME2 amplicon followed by BLAST analysis confirmed the predicted ORF, however a mismatch at 687 bp position was observed relative to the reference OU755558.1 genome assembly. To determine whether LmME2 is expressed, RT-PCR was performed using gene-specific primers (P1, P2), which confirmed the presence of LmME2 transcripts in *L. major* promastigotes (Fig. 1C). Expression in the intracellular amastigote stage was examined using an *in vitro* macrophage infection model. RT-PCR performed with LmME2-specific primers generated the expected amplicon from cDNA derived from *L. major* infected J774A.1 macrophages but not from uninfected controls (Fig. 1D), confirming the presence of LmME2 mRNA in intracellular amastigotes. Together, these findings establish that the second malic enzyme isoform, LmME2 that is expressed in both promastigote and amastigote stages of *L. major*.

**Fig. 1.**
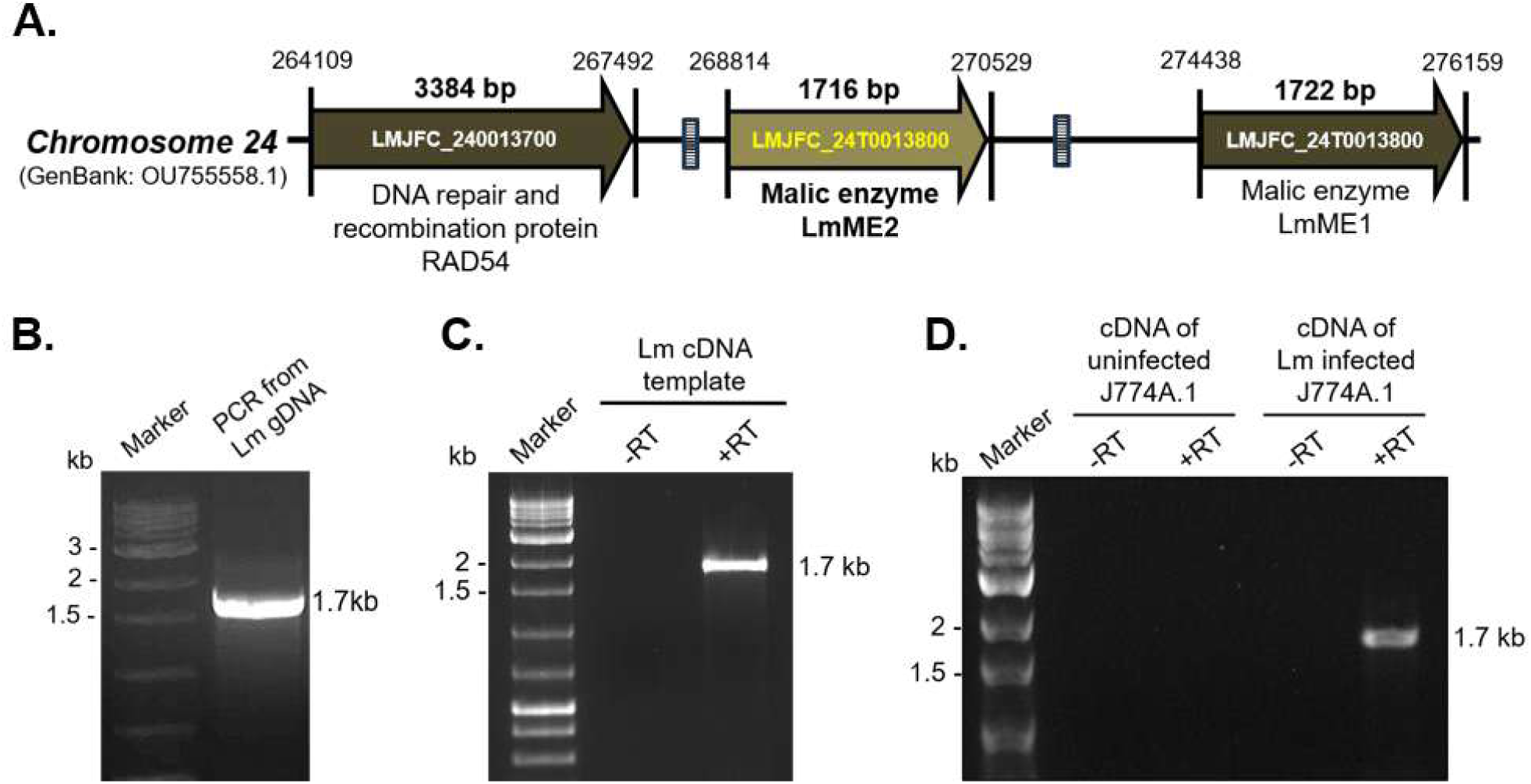
The second isoform of *L. major* malic enzyme, LmME2, is present and expressed in both promastigote and amastigote stages of *L. major*. (A) Schematic representation of the genomic locus of LmME2 (LMJFC_24T0013800) in the revised *L. major* genome assembly, highlighting that the two ME paralogues are located in tandem in the chromosome 24. The LmME2 locus is also flanked by SIDER elements, marked by small patterned rectangles. (B) Genomic DNA was isolated from wild-type *L. major* promastigotes and LmME2 (1716 bp) was amplified by full-length primers P1/P2. (C) Total RNA was isolated from wild-type *L. major* promastigotes and cDNA was synthesised followed by semi-quantitative RT-PCR amplification of LmME2 using primers P1/P2 (represented by lane marked as ‘+RT’). Respective negative control reaction (represented as ‘-RT’) was set up without reverse transcriptase (RT). (D) J774A.1 murine macrophages were either uninfected or infected with late stationary-phase *L. major* promastigotes (MOI 1:30). Infection was allowed for 12 hours after which uninternalized parasite were washed off and the cells were further incubated for 48 hours. Total RNA was isolated and cDNA was synthesised from both uninfected and infected macrophages. Semi-quantitative RT-PCR was performed with primers P1/P2. (represented by lanes marked as ‘+RT’). Respective negative control reactions (represented as ‘-RT’) were set up without RT.

### LmME2 is a NADP^+^-dependent malic enzyme regulated by oxaloacetate, ATP and fumarate

Following confirmation of LmME2 expression in *L. major*, we next examined whether it encodes a catalytically active malic enzyme capable of interconverting malate and pyruvate (Fig. 2A). The full-length LmME2 coding sequence was cloned into pET28a and expressed as an N-terminal His-tagged recombinant protein in BL21(DE3) *Escherichia coli*. The enzyme was purified to near homogeneity using Ni-NTA affinity chromatography (Fig. S1A) and subjected to enzymatic characterization. Purified LmME2 catalyzed both NADP^+^- dependent oxidative decarboxylation of malate and NADPH-dependent reductive carboxylation of pyruvate. The specific activities for malate decarboxylation and pyruvate carboxylation were ∼23 U/mg and ∼28 U/mg, respectively. Kinetic analysis revealed a lower Km for malate than for pyruvate, indicating a higher substrate affinity for malate (Table 1; Fig. S1B). Interestingly, while the specific activities of purified LmME2 were comparable, the whole-cell lysates of *L. major* displayed higher pyruvate carboxylation than malate decarboxylation activity (Fig. 2B). In this context it is important to note that the previously characterised LmME1 exhibited predominant malate decarboxylation activity, in its purified form (21). The shift in dominant reaction direction in the whole cell lysates suggested the presence of intracellular factors capable of modulating ME activity. Malic enzymes are known to be regulated by intermediary metabolites (39, 41, 42). We therefore tested key metabolic intermediates known to regulate malic enzymes in other systems also affect LmME2. Oxaloacetate, fumarate and ATP inhibited malate decarboxylation by 2.6-fold, 1.7-fold and 1.3-fold, respectively (Fig. 2C). In contrast, pyruvate carboxylation was selectively inhibited by ATP (1.9-fold), whereas oxaloacetate and fumarate had no significant effect (Fig. 2D), indicating differential regulation of the two catalytic directions of LmME2. To determine whether inhibition resulted from direct metabolite binding, intrinsic tryptophan fluorescence quenching assays were performed with purified LmME2 in the presence of increasing concentrations of each regulator. All three metabolites induced concentration-dependent fluorescence quenching, consistent with direct interaction with the enzyme (Fig. S1C, S1D, S1E). To understand the structural basis of metabolite-mediated regulation of LmME2 activity, AlphaFold3-based structural modelling of LmME2 was performed to generate a high-confidence dimer model with ipTM score of 0.84 and pTM score of 0.87, as no experimental structure is currently available for leishmanial malic enzymes. The binding sites for substrates (malate and pyruvate) along with cofactor NADP^+^ and coenzyme Mn^2+^ could be identified in the predicted LmME2 structure (Fig. S1F). Molecular docking analysis identified plausible binding sites for each regulator. Oxaloacetate exhibited a stable binding mode through hydrogen bonding with Glu53, Glu58 and Thr32, complemented by van der Waals interactions with Ile82, Thr85, Trp121, Val122 and Gly31 (chain A) (Fig. 2E). Fumarate was predicted to bind near the dimer interface, where the binding pocket is formed predominantly by residues from chain B with a contribution from His78 of chain A; it formed hydrogen bonds with Arg123 and Arg125, supported by hydrophobic contacts with Trp121, Val80, and Tyr120 (chain B) (Fig. 2E). On the other hand, ATP docked proximal to the NADP^+^-binding pocket, potentially explaining its inhibitory effect on both reaction directions. The binding site involved residues from chain A, with hydrogen bonds to Glu99, Asn463, Phe447 and Lys458, and additional stabilization was provided by van der Waals contacts involving Leu513, Gln461, Leu102, Pro103, Ala449, Pro448 and π-interactions with Glu99, Glu100 and Pro446, suggesting potential conformational modulation of ME2 activity (Fig. 2E). Taken together, these findings establish LmME2 as an active, bidirectional NADP^+^-dependent malic enzyme subject to metabolite-specific regulation, and identify plausible ligand-binding sites that may influence enzyme activity.

**Fig. 2.**
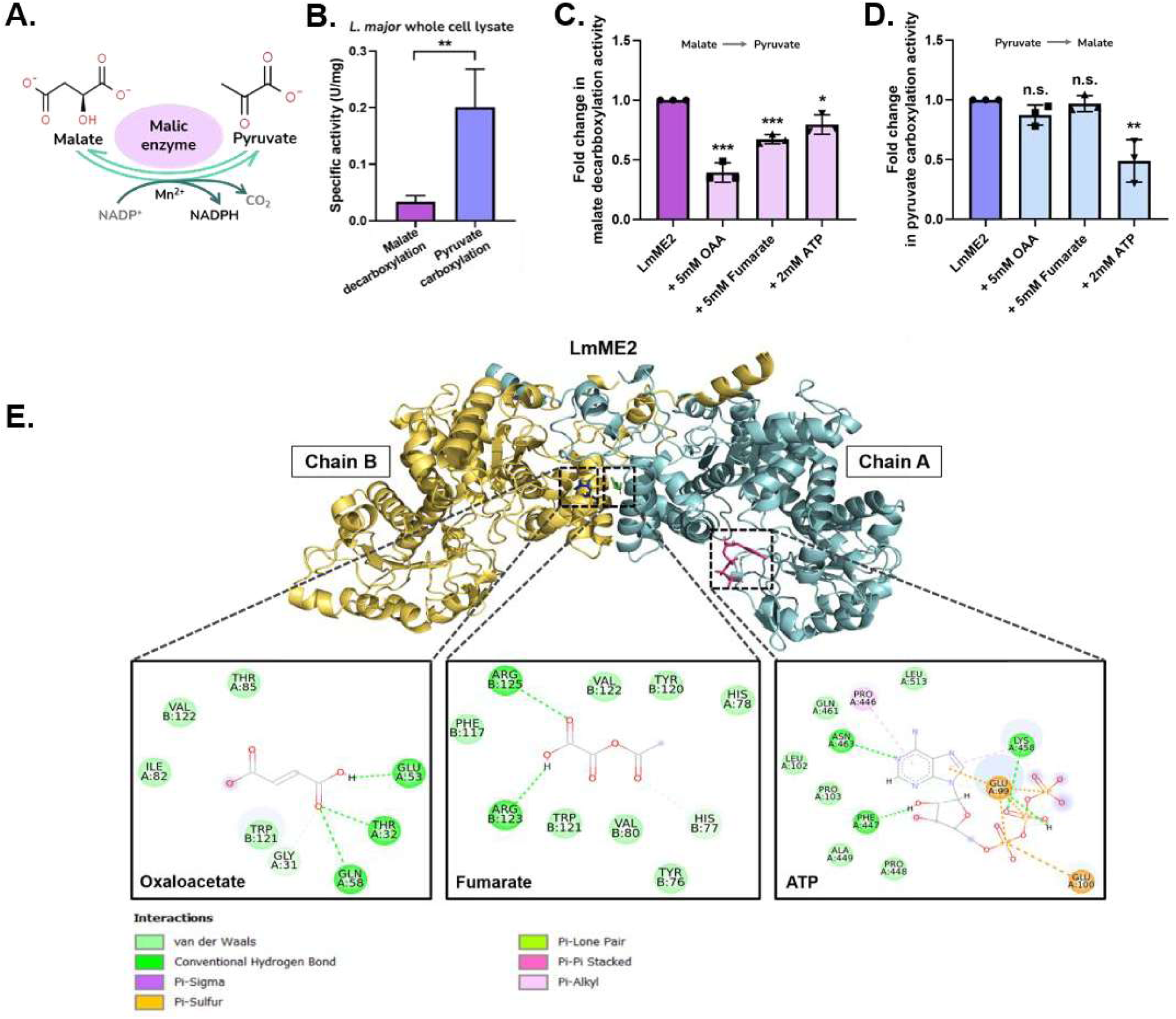
LmME2 activity is regulated by metabolites oxaloacetate, fumarate and ATP *in vitro* due to direct binding. (A) Schematic representation of the reversible reaction catalyzed by malic enzymes (MEs) using a divalent cation Mn^2+^ as cofactor, showing that NADP^+^ is reduced to generate NADPH during the decarboxylation reaction of malate to pyruvate. (B) Total malic enzyme activity was determined in wild-type *L. major* promastigotes with 20 µg whole cell lysate and specific activities were calculated. Bar graph for specific activities for malate decarboxylation and pyruvate carboxylation reactions. Error bars represent the mean±SD from 3 independent experiments. Statistical significance was tested by paired Student’s t-test. Asterisks indicate significant difference. Purified recombinant LmME2 was subjected to malic enzyme activity assay in absence or presence of regulatory metabolites (5mM oxaloacetate, 5mM fumarate, 2mM ATP). Bar graphs for fold change in (C) malate de- -carboxylation and (D) pyruvate carboxylation activities in presence of regulators with respect to no regulator added control. Error bars represent the mean±SD from 3 independent experiments. (E) 2D interactions of LmME2 with three regulators: oxaloacetate, fumarate and ATP. Oxaloacetate forms hydrogen bonding interactions with Glu53, Thr32, and Gln58, with additional van der Waals interactions with Ile82, Thr85, Trp121, Val122 and Gly31. Fumarate interacts with Arg125 and Arg123 at the dimer interface through hydrogen bonding, along with hydrophobic contacts with Trp121, Val80, and Tyr120. ATP forms hydrogen-bonding interactions with residues Lys458, Asn463, Phe447, and Glu99, with additional van der Waals interactions with Leu513, Gln461, Leu102, Pro103, Ala449, Pro448 and π-interactions with Glu99, Glu100 and Pro446. These interactions highlight the distinct binding modes of each regulator within LmME2. Statistical significance was tested by paired Student’s t-test. Asterisks indicate significant difference. *P ≤ 0.05, **P ≤ 0.01, ***P ≤ 0.001, n.s.- not significant.

**Table 1.** Kinetic parameters of purified LmME2. The kinetic parameters of LmME2 were determined by performing the malic enzyme activity assay with varying concentrations of substrates malate or pyruvate, whereas the concentrations of coenzymes NADP^+^ or NADPH as well as that of cofactor Mn^2+^ were kept constant. The apparent K_m_ and K_cat_ values were calculated by non-linear regression using SOLVER module in Microsoft Excel 2019. The parameters are the means of at least three independent calculations ± SD.

| Parameters | Malate decarboxylation | Pyruvate carboxylation |
| --- | --- | --- |
| **Specific activity (U/mg)** | 22.80 ± 2.51 | 27.92 ± 5.18 |
| **K<sub>m</sub> (mM)** | 0.25 ± 0.01 | 12.80 ± 2.45 |
| **K<sub>cat</sub> (s<sup>-1</sup>)** | 19.88 ± 2.94 | 39.72 ± 7.74 |

### LmME2 localizes to the cytosol in both promastigote and amastigote stages

Having established LmME2 as an active NADP^+^-dependent enzyme regulated by cellular metabolites, we next examined its subcellular localization. In silico analyses using TargetP and DeepLoc did not identify any organellar targeting sequence and predicted a cytosolic localization for LmME2. To experimentally determine its subcellular localisation, we raised polyclonal antibodies against LmME2 in mice and confirmed their isoform specificity by immunoblotting against purified recombinant LmME2 and LmME1 (Fig. S2A). The antibodies also detected endogenous LmME2 in *L. major* whole-cell lysate (Fig. S2B). We then subjected *L. major* promastigote lysate to sucrose density-gradient ultracentrifugation and performed immunoblotting of the fractions with anti-LmME2 antibody (Fig. 3A). The fractions were probed in parallel for phosphoglycerate mutase (PGAM), an established cytosolic marker in *Leishmania* (43). LmME2 co-fractionated with LmPGAM (Fig. 3B), indicating its cytosolic localization. To further confirm this finding, endogenous LmME2 was C-terminally tagged with mNeonGreen (mNG) using CRISPR-Cas9-mediated genome editing (Fig. S2C). The tagging was performed in the background of previously generated *L. major* strain stably expressing SpCas9 and T7 RNA polymerase (LmCas9/T7) (44, 45). sgRNA was designed such that the cleaved C-terminal region was repaired with a donor DNA cassette containing the mNG tag and a blasticidin-resistance gene (Fig. S2C). Successful tagging was verified by immunoblotting with anti-mNG antibody, which detected a ∼90 kDa band corresponding to the fusion of 63 kDa LmME2 with 28 kDa mNG (Fig. S2D). Fluorescence microscopy of LmME2-mNG promastigotes revealed a diffuse signal throughout the cell body, consistent with cytosolic distribution (Fig. 3C). Immunostaining of wild-type promastigotes with antibodies against LmME2 and LmPGAM showed clear co-localization and diffuse cytosolic staining (Fig. S2E), providing independent validation. As metabolic enzymes in trypanosomatids can undergo life stage-dependent relocalization (46), we next examined LmME2 localization in intracellular amastigotes. For that, J774A.1 macrophages were infected with late stationary-phase LmME2-mNG promastigotes. At 36 h post-infection, intracellular amastigotes displayed a similarly diffuse mNG signal throughout the parasite cytoplasm (Fig. 3D). Taken together, these results establish LmME2 as a cytosolic malic enzyme expressed in both the promastigote and amastigote life stages of *L. major*.

**Fig. 3.**
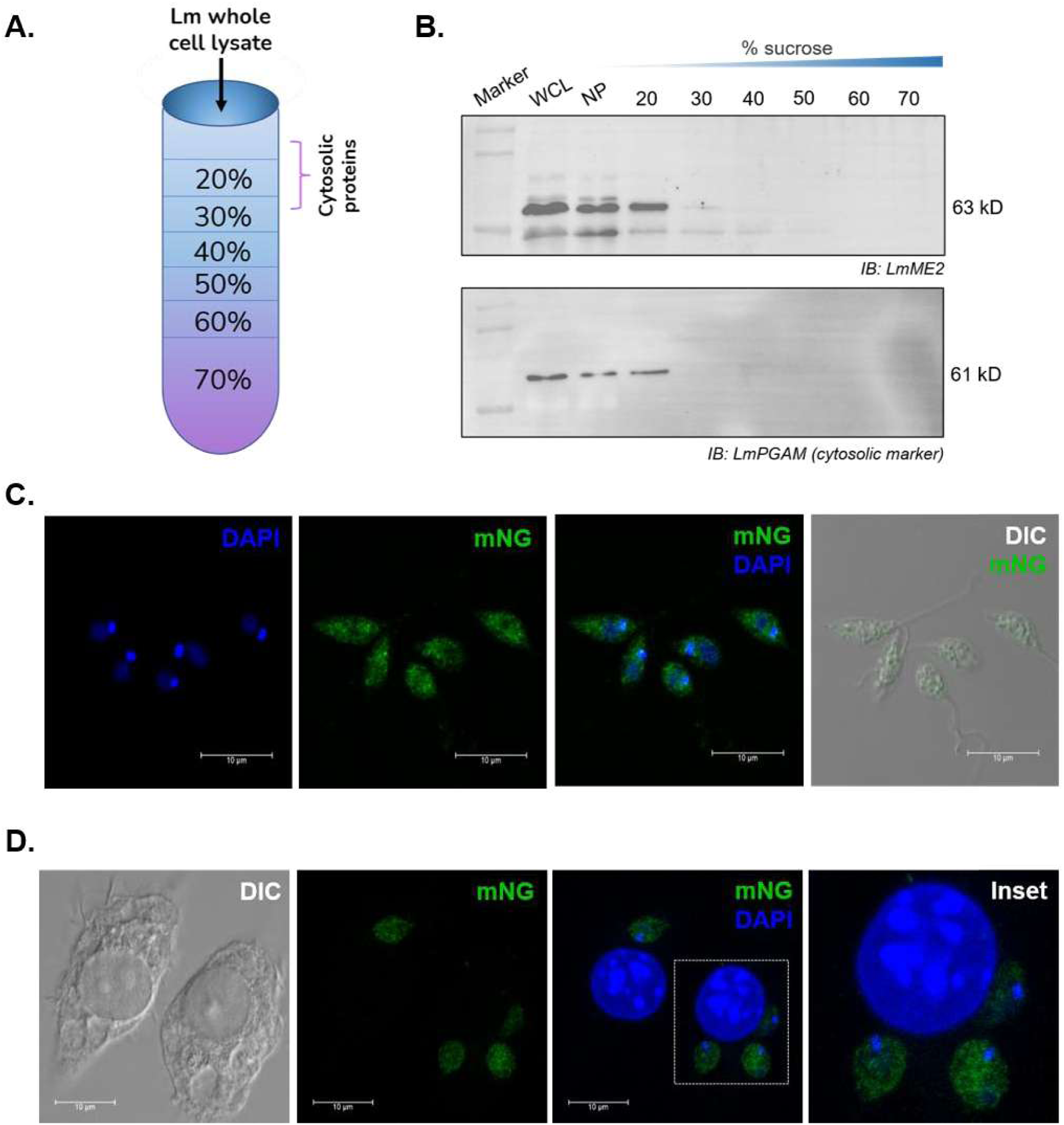
LmME2 is localised in the cytosol of *L. major* parasites. (A) Schematic representation of the sucrose density gradients used for subcellular fractionation by ultracentrifugation, showing cytosolic proteins to be settled in the topmost layer of 20% sucrose. (B) Wild-type *L. major* promastigotes were subjected to subcellular fractionation by sucrose-density gradient ultracentrifugation. Presence of LmME2 (62.7 kD) in the fractions were determined by western blotting with anti-LmME2 antibody. Purity of the cytosolic fraction was ensured by using antibody against LmPGAM (61 kD), a known cytosolic marker. (C) Representative images of cytosolic localization of LmME2 in *L. major* promastigotes. *L. major* promastigotes expressing LmME2-mNG (green) were fixed and mounted in DAPI (blue) containing mounting medium which stained both the nuclei and kinetoplast. The cells were visualised under Leica SP8 confocal microscope. Scale bars = 10 µm. (D) Representative images of localization of LmME2 in *L. major* amastigotes. J774A.1 macrophages were infected with *L. major* promastigotes expressing LmME2-mNG (green). 36 hours post infection, the cells were fixed and mounted in DAPI (blue) containing mounting medium. The cells were visualised under Leica SP8 confocal microscope. Scale bars = 10 µm.

### Deletion of LmME2, but not LmME1, disrupts pyridine nucleotide homeostasis in *L. major* promastigotes

To investigate the physiological role of LmME2, an LmME2 knockout (LmME2^-/-^) *L. major* strain was generated using CRISPR-Cas9-mediated genome editing in the LmCas9/T7 background. sgRNAs targeting the 5’ and 3’ UTRs of LmME2 were designed at the nearest PAM sites (6 bp upstream of the start codon and 7 bp downstream of the stop codon, respectively), to facilitate replacement of the LmME2 locus with puromycin-resistance cassette (Fig. 4A). Successful deletion of LmME2 was confirmed by immunoblotting of whole-cell lysates, which detected the protein in control parasites (LmCas9/T7) but not in LmME2^-/-^ parasites (Fig. 4B). Loss of LmME2 was further verified by immunofluorescence microscopy using anti-LmME2 antibodies, which showed the expected cytosolic signal in control parasites, but no detectable staining in LmME2^-/-^ promastigotes (Fig. S3A). Despite the absence of LmME2, LmME2^-/-^ promastigotes displayed no detectable morphological alterations, as examined by scanning electron microscopy (SEM) (Fig. S3B). Additionally, the growth rates of the knockout parasites remained comparable to control cells (Fig. S3C). To assess the contribution of LmME2 to total ME activity, ME assay was performed with whole cell lysates. Deletion of LmME2 resulted in a ∼60% reduction in malate decarboxylation activity and a ∼40% decrease in pyruvate carboxylation activity relative to control cells, indicating that LmME2 contributes substantially to both catalytic directions (Fig. S3D, S3E). Given that LmME2 catalyses an NADP^+^-dependent redox reaction, we next examined whether its deletion alters intracellular NADP^+^/NADPH homeostasis. Quantification of NADP^+^ and NADPH levels revealed that the NADPH/NADP^+^ ratio increased ∼1.7-fold in LmME2^-/-^ parasites relative to control cells (Fig. 4C). This increase was primarily driven by a ∼30% decrease in NADP^+^, while NADPH remained largely unchanged (Fig. 4D, 4E). Since NADP^+^ is generated from NAD^+^ via NAD^+^ kinase-mediated phosphorylation (Fig. 4F) (47, 48), we further examined the NAD(H) pool. The NADH/NAD^+^ ratio increased ∼1.8-fold in LmME2^-/-^ parasites (Fig. 4G), largely due to a ∼50% reduction in NAD^+^ levels, with minimal change in NADH levels (Fig. 4H, 4I). Together, these findings demonstrate that loss of LmME2 leads to depletion of both NADP^+^ and NAD^+^ pools. To determine whether this phenotype was specific to LmME2 or reflected a general consequence of malic enzyme deficiency, an LmME1 knockout strain was generated using a similar CRISPR-Cas9-mediated strategy. Deletion of LmME1 was confirmed by immunoblotting using anti-LmME1 antibodies (Fig. S4A). Kinetic assays revealed that LmME1 deletion caused a ∼80% reduction in malate decarboxylation activity and a ∼30% decrease in pyruvate carboxylation activity compared to control parasites (Fig. S4B, S4C). Interestingly, in contrast to LmME2 deletion, loss of LmME1 did not significantly alter either the NADPH/NADP^+^ or NADH/NAD^+^ ratios (Fig. S4D, S4E). Collectively, these findings identify LmME2, but not LmME1, as a major contributor to pyridine nucleotide homeostasis in *L. major* promastigotes.

**Fig. 4.**
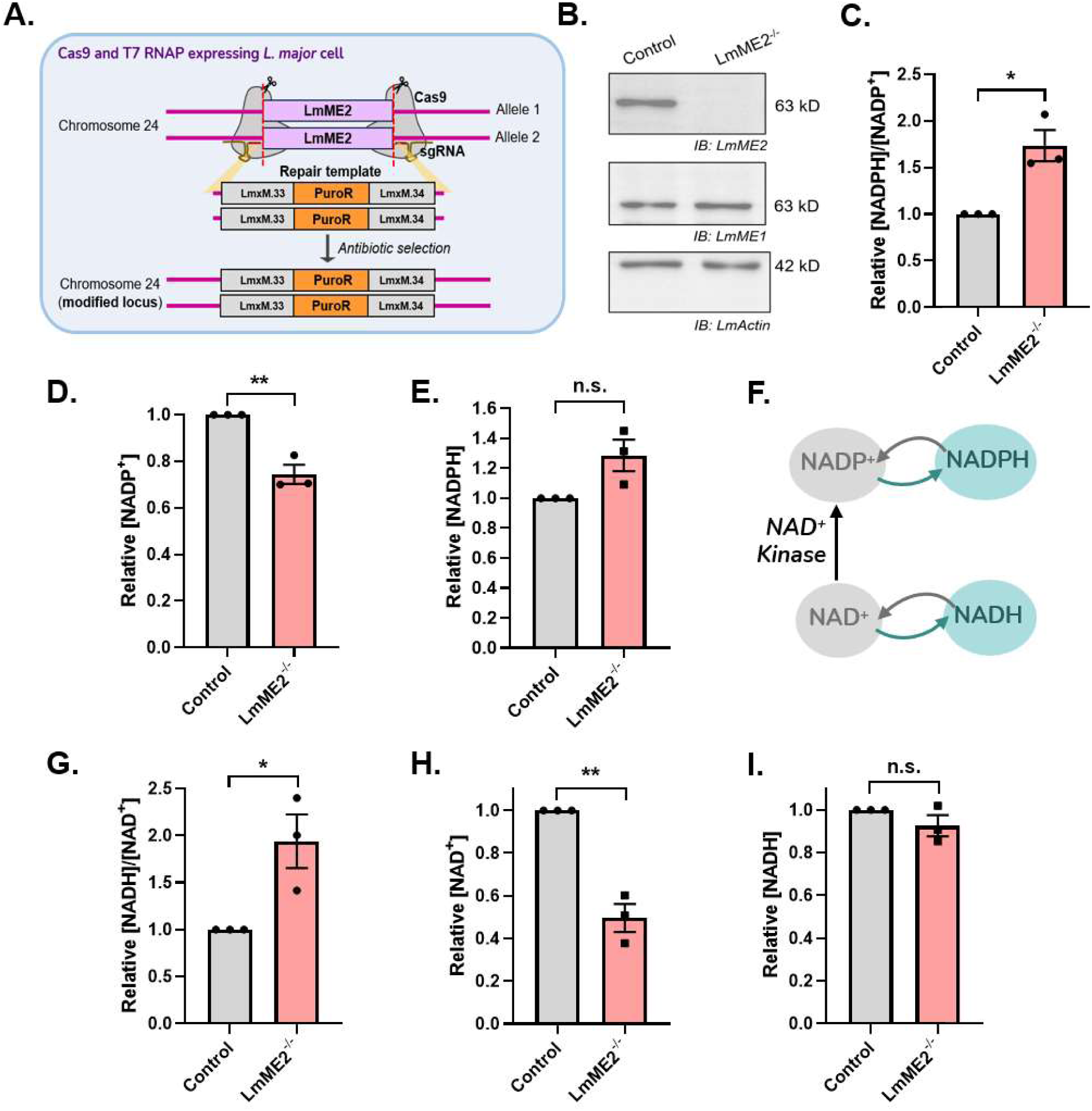
Deletion of LmME2 perturbs pyridine nucleotide homeostasis in *L. major* promastigotes. (A) Schematic representation of LmME2 locus in chromosome 24 of the *L. major* genome and the repair template containing the puromycin N-acetyltransferase (PuroR) gene flanked by LmxM.33 UTRs. Following electroporation of repair template along with sgDNA template into LmCas9/T7 strain, the sgDNA templates is *in vivo* transcribed to sgRNA, Cas9 mediated double strand break is repaired using the repair templates by homology-directed repair mechanism. (B) Post antibiotic selection, generation of LmME2^-/-^strain was verified by western blotting from whole cell lysates of control cells (LmCas9/T7) and LmME2^-/-^ strain with antibodies against LmME2. The presence of the other isoform, LmME1 was determined by western blotting with anti-LmME1 antibodies. LmActin was used as housekeeping control. NADPH and NADP^+^ levels were measured for late log phase *L. major* promastigotes of control and LmME2^-/-^ strains. Bar graphs for relative (C) NADPH/NADP^+^ ratio, (D) NADP^+^ and (E) NADPH levels with respect to control (LmCas9/T7) strain. Error bars represent mean±SEM. of values from 3 independent experiments. Statistical significance was tested by unpaired Student’s t- test. (F) Schematic representation of NADP+/NADPH pool and its replenishment from NAD^+^/NADH pool. NADP^+^ to NADPH turnover is maintained by several dehydrogenases and reductase enzymes, however, NADP^+^ cannot be de novo synthesised and can be replenished from NAD^+^ by NAD^+^ kinases. NAD^+^ to NADH turnover is again maintained by another set of dehydrogenases and reductases. NADH and NAD^+^ levels were also measured for late log phase *L. major* promastigotes of control (LmCas9/T7) and LmME2^-/-^ strains. Bar graphs for relative (G) NADH/NAD^+^ ratio (H) NAD^+^ and (I) NADH levels with respect to control (LmCas9/T7) strain. Error bars represent mean±SEM. of values from 3 independent experiments. Statistical significance was tested by unpaired Student’s t- test. Asterisks indicate significant difference. *P ≤ 0.05, **P ≤ 0.01, n.s. not significant

### LmME2 contributes to intracellular redox homeostasis and is critical for intracellular parasite survival and virulence in *L. major*

Despite perturbation of NAD(H) and NADP(H) pools, LmME2^-/-^ promastigote growth remained unaffected under standard culture conditions, suggesting that the contribution of LmME2 may become more pronounced under metabolically stringent and oxidative conditions of the macrophage phagolysosome. To model the oxidative stress encountered during infection, the parasites were treated with 350 μM H_2_O_2_ for 30 minutes. Intracellular ROS levels were increased in LmME2^-/-^ promastigotes by ∼1.2-fold under basal conditions and ∼1.4-fold following H_2_O_2_ treatment relative to control parasites (Fig. 5A). These findings indicate that loss of LmME2 compromises intracellular redox homeostasis, particularly under oxidative stress. Under glucose-replete conditions, the oxidative branch of the pentose phosphate pathway (PPP) serves as a major source of cytosolic NADPH required for antioxidant defence, with glucose-6-phosphate dehydrogenase (G6PD) catalysing its rate-limiting step (49, 50). However, under glucose-limiting conditions such as those encountered by intracellular amastigotes within the phagolysosome, flux through the PPP is expected to decrease, increasing reliance on parallel NADPH-generating pathways, including ME-mediated reduction of NADP^+^. To determine whether LmME2 becomes particularly important when PPP-derived NADPH production is restricted, parasites were treated with G6PDi, a pharmacological inhibitor of G6PD (51, 52). After 72 h of treatment with 4 μM G6PDi, LmME2^-/-^ parasites exhibited a ∼23% reduction in total cell number relative to control parasites (Fig. 5B). This heightened sensitivity to G6PD inhibition suggests that LmME2 helps sustain NADPH-dependent redox defence when PPP flux is limited, a function that may be particularly important for parasite survival within the glucose-restricted phagolysosomal environment. Given the role of LmME2 in maintaining redox balance and parasite fitness, we next assessed whether LmME2 deletion affected parasite virulence. J774A.1 murine macrophages infected with LmME2^-/-^ parasites displayed a ∼70% reduction in intracellular parasite burden compared with cells infected with control parasites (Fig. 5C), indicating impaired survival within host macrophages. This observation was further validated *in vivo* using the BALB/c mouse footpad infection model. Mice infected with control parasites developed progressively enlarging lesions that reached a lesion score of ∼1.5 mm^2^ by 12-week post infection, whereas mice infected with LmME2^-/-^ parasites exhibited minimal lesion development throughout the course of infection, with lesion scores remaining close to baseline and reaching only ∼0.01 mm² at the experimental endpoint (Fig. 5D, 5E). Although parasites remained detectable in the footpad homogenate from the LmME2^-/-^ infection group, the parasite load was approximately 100-fold lower than that observed in control infections (Fig. 5F). Collectively, these findings establish LmME2 as an important determinant of *L. major* virulence and support a model in which LmME2 promotes intracellular survival by maintaining pyridine nucleotide homeostasis and redox balance under metabolically restrictive conditions.

**Fig. 5.**
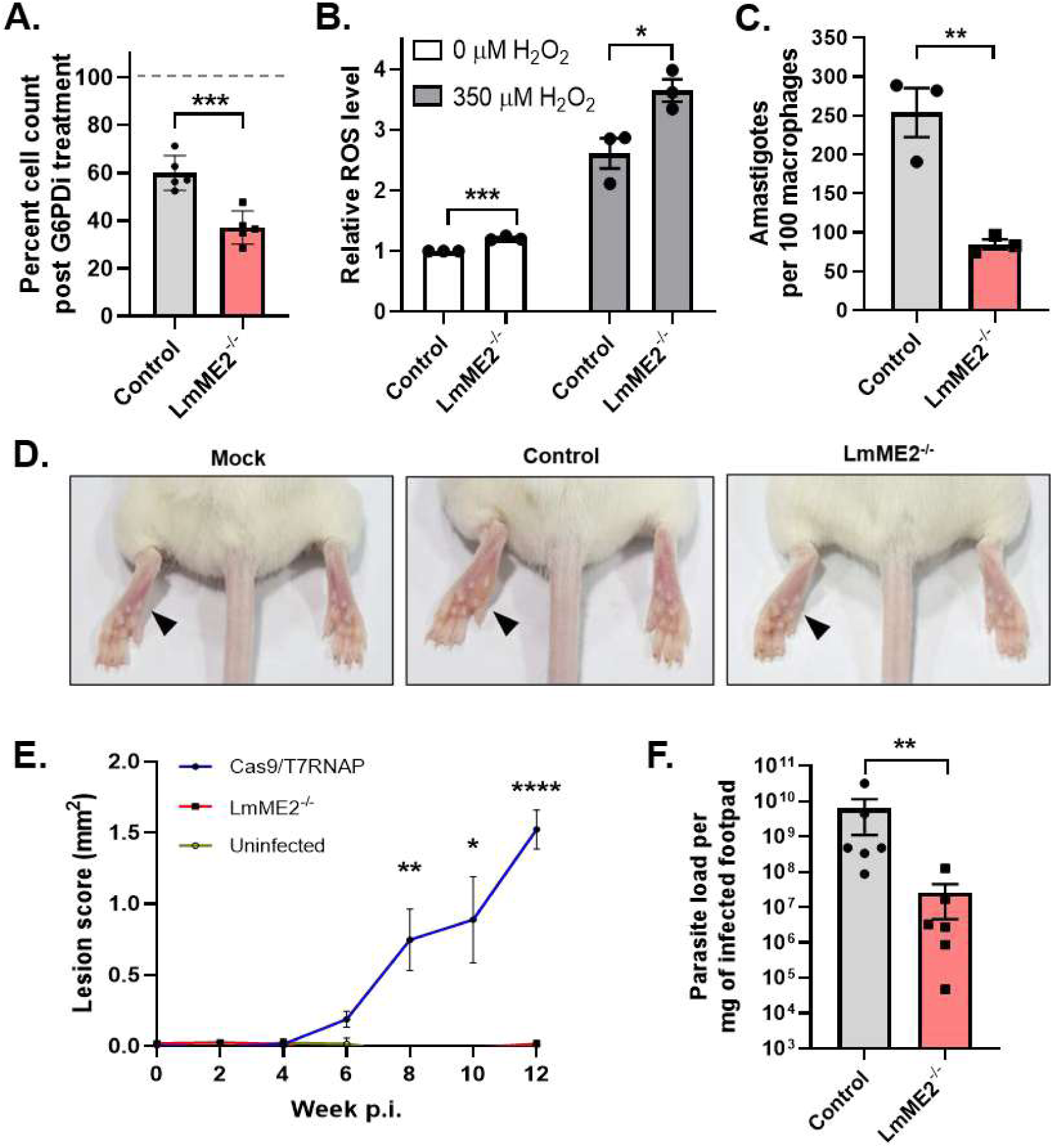
LmME2 deletion perturbs redox homeostasis in *L. major* parasites leading to drastically reduced parasite virulence. (A) *L. major* promastigotes of control (LmCas9/T7) and LmME2^-/-^ strains were grown in absence or presence of G6PDi, an inhibitor of the pentose phosphate pathway enzyme glucose-6-phosphate dehydrogenase (G6PDH) for 72 h. Bar graphs represent percent cell count of control (grey) and LmME2^-/-^ (red) strains post treatment with 4 µM G6PDi relative to cell count of untreated control strain. Error bars represent mean±SEM. of values from 3 independent experiments. (B) Intracellular ROS for control and LmME2^-/-^ strains at both basal and oxidative stress induced conditions were measured by CM-H_2_DCFDA assay. Oxidative stress was induced by 350 µM H_2_O_2_ for 30 minutes. Bar graphs represent relative ROS level of LmME2^-/-^ strain with respect to control (LmCas9/T7) strain at both basal (white bars) and H_2_O_2_ treated (grey bars) conditions. Error bars represent mean±SEM. of values from 3 independent experiments. Statistical significance was tested by unpaired Student’s t- test. (C) J774A.1 macrophages were infected with control (LmCas9/T7) (grey bar), LmME2^-/-^ (red bar) *L. major* promastigotes of late stationary phase. The cells were fixed 30 hrs. p.i., mounted in DAPI containing medium and imaged under Olympus IX83 epifluorescence microscope. From the images, the total number of DAPI-stained nuclei and kinetoplast duo of *Leishmania* amastigotes within macrophages in a field were counted to determine the intracellular parasite burden per 100 macrophages. Error bars represent mean±SEM of values from 3 independent experiments. Statistical significance was tested by unpaired Student’s t-test. BALB/c mice were either uninfected (mock) or infected with 5 x 10^6^ late stationary phase *L. major* promastigotes of control or LmME2^-/-^ strains by subcutaneous injection in their left hind footpad. (D) Representative images of the left hind footpads of uninfected and infected BALB/c mice at week 12 p.i., when the experiment was terminated. (E) Lesion scores were calculated from weekly measurements of width and thickness of the lesions of the infected footpads till week 12 p.i. (n=7 mice per set for 2 independent sets). Error bars represent mean±SEM. of values from 2 independent experiments. Statistical significance was tested by unpaired Student’s t-test. (F) Parasite burden in mice footpads infected with control (LmCas9/T7) (grey bar) and LmME2^-/-^ (red bar) parasites were determined by limiting dilution assay with the lysates of excised lesion tissues. (n=7 mice per set for 2 independent sets) Error bars represent mean±SEM. of values from 2 independent experiments. Statistical significance was tested by Mann-Whitney test. Asterisks indicate significant difference. *P ≤ 0.05, **P ≤ 0.01, ***P ≤ 0.001, n.s. not significant.

## Discussion

In the original *L. major* Friedlin genome assembly (FR796420.1 / NC_007265.2) (53), only a single malic enzyme was annotated, in contrast to the other *Leishmania* species, which were predicted to encode two ME isoforms. This led to the speculation that *L. major* may have lost the second isoform during evolution (39). However, the revised *L. major* genome assembly now contains two annotated ME paralogues, resolving this apparent misconception of the missing ME isoform. Interestingly, the LmME2 locus is flanked by SIDER elements, repetitive sequences known to be sites of assembly collapse in short-read sequencing of trypanosomatid genomes (54, 55); this likely accounts for the original miss in the initial genome assembly. The two paralogues share ∼65% amino acid sequence identity and lie ∼3.3 kb on chromosome 24, consistent with a tandem-duplication. Consistent with this interpretation, Alonso et al. identified several regions that were absent from the original genome assembly, one of which corresponds to the genomic locus containing LmME2 (55).

The present study, together with our previous characterisation of LmME1, establishes that *L. major* expresses two spatially distinct malic enzyme isoforms: a mitochondrial enzyme (LmME1) and a cytosolic enzyme (LmME2). The compartmentalisation of these enzymes suggests functional specialization within different metabolic environments of the parasite. While LmME1 has previously been shown to support gluconeogenesis through its pyruvate-carboxylating activity (21), the physiological significance of a cytosolic malic enzyme in *Leishmania* remained unknown. Our findings now demonstrate that LmME2 contributes to pyridine nucleotide homeostasis, redox balance, and parasite virulence, thereby providing the first experimental evidence for a functional cytosolic malic enzyme in the genus.

Biochemically, recombinant LmME2 functions as a bidirectional NADP^+^-dependent enzyme capable of interconverting malate and pyruvate. Although kinetic profiling revealed a lower Km for malate than for pyruvate in both LmME2 and LmME1, indicating higher substrate affinity in the decarboxylation direction, whole-cell *L. major* lysates predominantly exhibited pyruvate carboxylation activity. A similar preference was observed previously for LmME1, despite the purified enzyme favouring malate decarboxylation (21), indicating that the reaction direction is influenced by intracellular metabolic factors. Malic enzyme are widely regulated by post-translational modifications and intermediary metabolites (39, 56–59). Likewise, we found LmME2 to be regulated by key metabolites: ATP, fumarate and oxaloacetate. ATP inhibited both catalytic directions, whereas fumarate and oxaloacetate selectively inhibited malate decarboxylation through distinct binding sites predicted by structural modelling and docking analyses. Notably, this regulatory profile differs from that of mammalian malic enzymes, where fumarate typically acts as an activator (41, 60). This contrasting regulatory response highlights a fundamental divergence in metabolic control between host and parasite enzymes, suggesting that LmME2 may be selectively targeted without affecting the host orthologue.

A central finding of this study is the role of LmME2 in maintaining intracellular pyridine nucleotide homeostasis. Cytosolic malic enzymes are generally viewed as sources of NADPH that support reductive biosynthesis and antioxidant defence. Surprisingly, deletion of LmME2 did not significantly alter NADPH levels, suggesting compensation by alternative NADPH-generating pathways operating in promastigotes, including the oxidative pentose phosphate pathway. Notably, no significant transcriptional upregulation of the pentose phosphate pathway or trypanothione biosynthesis enzymes was observed, indicating that LmME2 deletion does not trigger a detectable compensatory response at the transcript level.

Strikingly, the most prominent phenotype was a marked depletion of the oxidised pyridine nucleotide pools, with NAD^+^ and NADP^+^ levels reduced by approximately 50% and 30%, respectively. Given that most studies examining perturbation of malic enzymes report effects primarily on NADPH (26, 29) with relatively limited impact on NAD^+^, our findings point to a previously unrecognised role of malic enzyme in maintaining global pyridine nucleotide balance. The underlying mechanism remains unclear and may involve altered de novo NAD^+^ biosynthesis, salvage pathway flux, or changes in NAD^+^ consuming reactions such as sirtuin or ADP-ribosyltransferase-mediated processes (61). Importantly, this phenotype was specific to LmME2, as deletion of the mitochondrial isoform LmME1 produced no comparable changes, highlighting the distinct physiological functions of the two enzymes despite their shared catalytic activity.

The consequences of this metabolic defect became particularly evident under conditions that mimic the intracellular amastigote environment. Within the macrophage phagolysosome, parasites experience sustained oxidative stress together with restricted glucose availability, conditions expected to limit flux through the pentose phosphate pathway and increase reliance on alternative NADPH-generating reactions. Consistent with this model, LmME2-deficient parasites accumulated higher levels of reactive oxygen species and displayed increased sensitivity to pharmacological inhibition of glucose-6-phosphate dehydrogenase, directly demonstrating the increased dependence on LmME2-derived reducing cofactors in the limiting PPP flux condition. However, despite the loss of LmME2 and the substantial depletion of NAD^+^ and NADP^+^, the corresponding reduced nucleotides, NADH and NADPH, remained largely unchanged. This pattern suggests that the parasite prioritises maintenance of the reduced pyridine nucleotide pools, potentially through compensatory metabolic mechanisms, even at the expense of the oxidised nucleotide pools. Whether such compensatory responses contribute directly to the depletion of NAD^+^ and NADP^+^ observed in the knockout parasites remains an important question for future investigation.

The physiological importance of LmME2 was eventually reflected in the pronounced virulence defects of the knockout parasites, which exhibited markedly reduced survival within macrophages and a ∼100-fold reduction in parasite burden in infected mice. Collectively, our findings identify LmME2 as a previously unrecognized regulator of pyridine nucleotide homeostasis and redox adaptation in *L. major*. Beyond establishing the existence and function of a cytosolic malic enzyme in *Leishmania*, this work reveals a non-canonical role for malic enzyme in maintaining NAD^+^ and NADP^+^ pools and demonstrates the importance of this function for parasite survival under host-imposed stress. These findings broaden our understanding of metabolic adaptation in trypanosomatid parasites and highlight LmME2 as a potential target for antileishmanial drug development.

## Materials and methods

Unless otherwise mentioned, all reagents were purchased from Sigma-Aldrich. All primers were procured from Integrated DNA Technologies and their sequence details are provided in Table S1. Details for the antibodies used and their dilutions are provided in Table S2.

### *Leishmania* parasite and mammalian cell culture

*Leishmania major* strain (MHOM/SU/73/5-ASKH) and murine macrophage cell line J774A.1 (ATCC TIB-67) were procured from American Type Culture Collection (ATCC). *L. major* promastigotes were cultured as previously described (62). Briefly, the parasites were grown at 26 °C in M199 medium (Gibco) supplemented with 15% heat-inactivated foetal bovine serum (FBS; Gibco), 23.5 mM HEPES, 0.2 mM adenine, 150 µg/ml folic acid, 10 µg/ml hemin, 120 U/ml penicillin, 120 µg/ml streptomycin, and 60 µg/ml gentamicin and pH was adjusted to 7.2. J774A.1 macrophages were cultured in Dulbecco’s Modified Eagle’s Medium (DMEM; Gibco), pH 7.4, supplemented with 10% heat-inactivated FBS, 2 mM L-glutamine, 100 μg/ml streptomycin, and 100 U/ml penicillin. The cells were maintained at 37 °C in a humidified incubator with 5% CO_2_. Cell count was determined, where required, using a hemacytometer following fixation with 0.9% formal saline or staining with Trypan blue to assess cell viability. For all experiments, antibiotic selection was omitted from the culture medium unless otherwise mentioned.

### Genomic analysis & PCR

Genomic DNA was isolated from *L. major* promastigotes using the phenol-chloroform extraction method. Briefly, ∼1 × 10□ cells were harvested by centrifugation at 1000 × g and resuspended in 100 µL lysis buffer (0.1 M Tris-HCl, 0.1 M NaCl, 1% SDS, 100 mM EDTA, 100 µg/mL Proteinase K). The suspension was incubated at 55 °C for 2 h to ensure complete cell lysis and protein digestion. Following lysis, an equal volume of phenol and chloroform (25 µL each) was added, and the mixture was centrifuged at 13,000 × g for 15 min. The upper aqueous phase was carefully transferred to a fresh tube and to which an equal volume of chloroform was added, followed by centrifugation at 13,000 × g for 15 min. The supernatant was collected, and 1/10 volume of 3 M sodium acetate and 2 volumes of absolute ethanol were added, followed by centrifugation at 13,000 × g for 10 min. The obtained pellet was washed twice with 70% ethanol, air-dried, and resuspended in nuclease-free water. DNA concentration and purity were determined spectrophotometrically using a NanoDrop instrument (Thermo Fisher Scientific). PCR amplification of genomic DNA was performed using Leo Mastermix (Dx/dt) and gene-specific primers according to the manufacturer’s protocol.

### Gene expression analysis in *L. major* promastigote

Total RNA was isolated from *L. major* promastigotes using TRIzol reagent (Invitrogen) according to the manufacturer’s instructions. The isolated RNA was treated with DNase I (Invitrogen) to remove residual genomic DNA contamination. cDNA was synthesized from DNase I-treated RNA using the Verso cDNA synthesis kit (Thermo Fisher Scientific) following the manufacturer’s protocol. Negative control reactions (-RT) were prepared in parallel without the reverse transcriptase enzyme, while including all other components. PCR amplification was performed using cDNA from both +RT and -RT reactions as templates.

### Gene expression analysis in *L. major* amastigote

Infection of J774A.1 macrophages with *L. major* was performed as described previously by us (62), with minor modifications. Briefly, macrophages were activated with 100 ng/mL lipopolysaccharide (LPS) for 6 h prior to infection. Stationary-phase *L. major* promastigotes were then added at a multiplicity of infection (MOI) of 30:1 (parasite:macrophage ratio). After 12 h, uninternalized parasites were removed by washing with phosphate-buffered saline (PBS), and fresh Dulbecco’s Modified Eagle’s Medium (DMEM) was added. Infected cells were further incubated for approximately 60 h. Total RNA was isolated from both infected and uninfected macrophages, and RT-PCR analysis was performed as described above.

### Purification of recombinant LmME2

The open reading frame (ORF) of the LmME2 gene was PCR-amplified from wild-type *L. major* genomic DNA using gene-specific primers (P3, P4). The amplified 1.7 kb full-length LmME2 was digested with EcoRI and NotI and cloned into the corresponding sites of the pET-28a(+) expression vector to generate an N-terminal 6×His-tagged construct. The integrity of the construct was confirmed by DNA sequencing. For recombinant protein expression, the plasmid was transformed into *Escherichia coli* BL21(DE3) cells. Transformed cells were cultured overnight in 5 mL Luria-Bertani (LB) medium containing 50 µg/mL kanamycin at 37 °C. The overnight culture was used to inoculate 500 mL LB medium, and cells were grown until an optical density at 600 nm of ∼0.6 was reached. Protein expression was induced with 0.5 mM isopropyl β-D-1-thiogalactopyranoside (IPTG), followed by incubation for 8 h at 20 °C. Cells were harvested by centrifugation at 3,000 × g for 10 mins and cell pellet was stored at -80 °C overnight. The cell pellet was resuspended in ice-cold lysis buffer (50 mM Tris-HCl, 100 mM NaCl, 20 mM imidazole, 0.5 mM DTT, 0.5% (v/v) Triton X-100, 5% (v/v) glycerol, 1 mg/mL lysozyme, and 1 mM phenylmethylsulfonyl fluoride (PMSF), pH 8.0). The suspension was incubated on ice for 30 min with intermittent mixing and subsequently lysed by French press (15kpsi pressure for three strokes). The lysate was clarified by centrifugation at 14,000 × g for 30 min at 4 °C. The supernatant was next incubated with Ni²c-nitrilotriacetic acid (Ni-NTA) agarose resin (Qiagen) for 1 h at 4 °C in an end-to-end shaker. The unbound proteins were separated by centrifugation at 2,000 × g for 3 min. The resin was washed sequentially with wash buffer (50 mM Tris-HCl, 300 mM NaCl, 30 mM imidazole, 1 mM PMSF, pH 8.0) and a second wash containing 40 mM imidazole. Bound His-tagged LmME2 protein was eluted using the same buffer supplemented with 250 mM imidazole, 0.5 mM DTT and 1% (v/v) glycerol. The eluted protein was dialyzed three times against dialysis buffer (50 mM Tris-HCl, 100 mM NaCl, 0.5 mM DTT, 1 mM PMSF, pH 8.0). Protein purity was assessed by 12% SDS-PAGE followed by Coomassie Brilliant Blue staining.

### Malic enzyme activity assay

Malic enzyme activity in purified LmME2 or *L. major* whole-cell lysates was measured spectrophotometrically using a U2900 spectrophotometer (Hitachi) with a 1 cm path length quartz cuvette, as described previously (21, 63) with minor modifications. For assaying malate decarboxylation, the reaction mixture (1 mL) contained malate buffer (50 mM Tris- HCl (pH 7.5), 10 mM malate, 1 mM MnCl_2_), and 0.15 mM NADP^+^. For pyruvate carboxylation, the assay mixture consisted of pyruvate buffer (50 mM Tris-HCl (pH 7.5), 50 mM pyruvate, 75 mM NaHCO_2_, 1 mM MnCl_2_), and 0.15 mM NADPH. Reaction mixtures were equilibrated at room temperature and prepared freshly prior to the assay. The reaction was initiated by addition of 4 µg purified enzyme or 20 µg whole-cell lysate. The change in absorbance at 340 nm was monitored over 2 min. Enzyme activity was calculated using the molar extinction coefficient of NADPH (6.22 mM^−1^·cm^−1^) and expressed as enzyme units (EU) per mg protein, where one unit corresponds to the production or consumption of 1 µmol NADPH per min. Protein concentration was determined by the method of Lowry et al. (64). For kinetic analysis, purified LmME2 was assayed across increasing concentrations of malate or pyruvate while maintaining other components at saturating levels. Initial reaction velocities were determined and fitted to the Michaelis-Menten equation using nonlinear regression analysis in Microsoft Excel. The turnover number (k_cat_) was calculated based on the molecular mass of the LmME2 monomer (62.8 kDa). All kinetic parameters represent the mean of at least three independent experiments.

### Enzyme activity regulation and regulator interaction analysis of LmME2

The effect of metabolic regulators on LmME2 activity was assessed using standard malic enzyme assay setup as described above. Briefly, purified recombinant LmME2 was assayed in absence or presence of regulatory metabolites oxaloacetate (5 mM), fumarate (5 mM), ATP (2 mM), and glucose (10 mM). Regulator ligands were freshly prepared by dissolving oxaloacetate in autoclaved double-distilled water, fumarate in DMSO, and ATP in 1 M Tris-HCl buffer (pH 7.5). To evaluate protein-ligand interactions, fluorescence quenching assays were performed using a spectrofluorometer (Hitachi F-7000) equipped with a xenon lamp. Intrinsic tryptophan fluorescence of LmME2 was monitored by exciting the protein at 295 nm, and emission spectra were recorded between 311 and 450 nm using a slit width of 5 nm and an integration time of 2 s. Increasing concentrations of each ligand were titrated against the protein, and changes in fluorescence intensity were used to assess ligand binding.

### Molecular docking analysis of LmME2 with metabolic regulators

Molecular docking studies were performed to investigate interactions between LmME2 and potential metabolic regulators, including ATP, oxaloacetate, and fumarate. Docking simulations were carried out using AutoDock 4.2. The LmME2 protein structure was prepared using the WHAT IF server, followed by conversion to PDBQT format using AutoDock tools. Ligand structures (ATP, oxaloacetate, and fumarate) were similarly prepared and converted to PDBQT format. During preparation, missing atoms were added, Kollman charges were assigned to the protein, and Gasteiger charges were assigned to the ligands. Docking simulations were performed with 50 independent runs, a population size of 300, and default energy evaluation parameters to ensure adequate conformational sampling. Docked conformations were ranked based on binding energy, and the lowest-energy poses were selected for further analysis. Protein-ligand interactions were visualized using Discovery Studio, and two-dimensional interaction maps were generated to characterize binding modes and key interacting residues.

### Polyclonal antibody generation against LmME2

Polyclonal antibodies against LmME2 were generated in BALB/c mice using a standard immunization protocol. Female mice (4-6 weeks old) were acclimatized for 1-2 weeks prior to immunization under institutional animal facility conditions. Recombinant LmME2 protein was purified prior to immunization as described above. A pre-immune blood sample was collected before antigen administration. For primary immunization (Day 0), mice were injected with 50 µg of purified recombinant LmME2 mixed with equal volume of Complete Freund’s Adjuvant (CFA; Santa Cruz Biotechnology). Booster immunizations were administered using 50 µg antigen emulsified in Incomplete Freund’s Adjuvant (IFA) at 2-week intervals for a total of three boosters. Seven days after the final booster injection, blood was collected for serum preparation. Blood samples were allowed to clot at room temperature and centrifuged at 10,000 × g for 10 min to separate serum containing polyclonal antibodies against LmME2, which was aliquoted and stored at -80 °C until further use.

### Western blotting

Whole-cell lysates of *L. major* promastigotes were prepared by resuspending cells in PBS supplemented with protease inhibitor cocktail, followed by sonication. Protein concentration was determined using the method Lowry et al (64). Equal amounts of protein were loaded onto a 12% SDS-polyacrylamide gel and transferred onto methanol-activated polyvinylidene difluoride (PVDF) membrane. The membrane was blocked with 5% (w/v) skim milk in TBST (10 mM Tris-HCl, 150 mM NaCl, 0.025% Tween-20) and washed twice with TBST for 5 min each. The membrane was incubated overnight at 4 °C with primary antibodies (mouse anti-LmME2, rabbit anti-mNeonGreen, rabbit anti-LmME1 or rabbit anti-LdActin; Table Sx) diluted in TBST. Following incubation, the membrane was washed five times with TBST (5 min each) and then incubated with appropriate horseradish peroxidase (HRP)-conjugated anti-rabbit or anti-mouse secondary antibody for 1.5 h at room temperature. After washing five times with TBST, protein bands were detected using a chemiluminescent substrate (Bio-Rad) and visualized using a ChemiDoc imaging system (Bio-Rad).

### Subcellular fractionation

Subcellular fractionation of *L. major* promastigotes was performed as described previously (65), with minor modifications. Briefly, ∼1 × 10c logarithmic-phase promastigotes were harvested by centrifugation at 1,000 × g for 5 min at 4 °C and washed twice with PBS to remove residual serum components. The cell pellet was resuspended in 500 µL hypotonic buffer (5 mM HEPES, 2 mM EGTA, 2 mM DTT) and lysed by mechanical shearing using 20 passes through a 31-gauge needle. The lysate was made isotonic by addition of 0.25 volumes of isotonic buffer (50 mM HEPES, 250 mM sucrose, 1 mM ATP, 1 mM EGTA) and centrifuged at 3,000 × g for 15 min at 4 °C to remove unlysed cells and nuclei. The resulting post-nuclear supernatant was layered on a 7 mL 20-60% sucrose gradient with 70% sucrose cushion and subjected to ultracentrifugation at 39,000 rpm for 6 h at 4 °C using an SW41Ti rotor in a Beckman L-80 ultracentrifuge (Beckman Coulter). Fractions were carefully collected from the top and transferred to separate tubes. Proteins from each fraction were concentrated using chloroform-methanol precipitation. Briefly, four volumes of methanol were added and mixed, followed by one volume of chloroform and three volumes of water with thorough vortexing after each addition. Samples were centrifuged at 14,000 × g for 30 min at 4 °C to achieve phase separation, with proteins localized at the interphase. The upper aqueous/methanol layer was removed without disturbing the interphase. Subsequently, 4.5 volumes of methanol were added, and samples were centrifuged at 14,000 × g for 15 min to pellet proteins. The proteins were precipitated along the walls of the tubes. The supernatant was discarded, and protein pellets were air-dried and resuspended in 200 µL lysis buffer. Protein concentration was determined using standard methods, and equal amounts of protein (30 µg) from each fraction were subjected to western blot analysis.

### Immunofluorescence studies

Glass coverslips were sterilized by immersion in 70% ethanol for 30 min under UV light, followed by flame drying. Sterile coverslips were placed in a 6-well plate and coated with poly-L-lysine (150 µL per coverslip) for 30 min to facilitate cell attachment. The poly-L-lysine was blotted off and ∼1 × 10c mid-log phase cells *L. major* promastigotes were then added to the coverslips and allowed to adhere for 30 min. on-adhered cells were removed by washing twice with PBS for ∼2-3 min each with gentle agitation. Cells were fixed with 1:1 acetone-methanol for 10 min at room temperature in the dark, followed by two washes with PBS. Permeabilization was performed using 0.1% Triton X-100 for 2 min, and coverslips were washed again with PBS. Blocking was carried out using 0.2% gelatin in PBS for 10 min. Coverslips were next incubated with primary antibodies diluted in blocking solution for 1.5 h, followed by two washes with PBS for 10 min each. Subsequently, samples were incubated with appropriate secondary antibodies (diluted in blocking buffer) for 2 h in the dark. Coverslips were washed twice with PBS of 10 min each. Excess PBS was removed, and coverslips were air-dried before mounting onto glass slides using DAPI-containing mounting medium. Slides were sealed and imaged using a confocal laser scanning microscope (Leica Microsystems SP8).

### CRISPR-Cas9 mediated C-terminal tagging and gene knockout

Genetic manipulation of the LmME2 or LmME1 locus in *L. major* was performed using a CRISPR-Cas9 system following the protocol reported by Beneke et al. (66). and as schematically represented in Fig. S3C, 4A. C-terminally tagged LmME2 (LmME2-mNG) and LmME2 or LmME1 knockout (LmME2^-/-^, LmME1^-/-^) strains were generated using *L. major* parasites constitutively expressing SpCas9 and T7 RNA polymerase (LmCas9/T7), as generated and previously (44, 45). Single guide RNA (sgRNA) templates or sgDNAs were designed manually for LmME2 or using the LeishGEdit platform for LmME1. The resulting sgRNAs were meant to either the 3′ end of the respective open reading frames (for C-terminal tagging) or sequences flanking the 5′ and 3′ untranslated regions (UTRs) for gene deletion. sgDNAs were generated by overlap extension PCR using gene-specific primers containing the T7 promoter and 20-nucleotide target sequence (P5/P6/P7/P12/P13), along with the universal G00 primer encoding the sgRNA scaffold. Donor DNA cassettes were PCR-amplified using primers incorporating 30-nucleotide homology arms corresponding to sequences flanking the Cas9 cleavage sites. For C-terminal tagging, the donor cassette (amplified from pPLOTv1 blast-mNeonGreen-blast plasmid, using primers P10, P11) encoded an in-frame fusion of mNeonGreen (mNG) at the 3′ end of LmME2 along with a blasticidin resistance marker, whereas for gene knockout, puromycin resistance cassette (amplified from pTPuro_v1 plasmid, using primers P8, P9 or P14, P15) was used to replace the endogenous LmME2 of LmME1 alleles. All PCR reactions were performed using high-fidelity Phusion polymerase (ThermoScientific) keeping the parameters similar as reported by Beneke et al. (66), and amplified donor fragments were purified by gel extraction (Bio Bharati Life Science). Purified sgDNAs and corresponding donor DNA fragments were co-electroporated into mid-log phase LmCas9/T7 promastigotes using BTX Gemini electroporator (480 V, 550 lF, 25 ohms) and selected under appropriate antibiotic pressure (5 µg/mL blasticidin S for C-terminal tagging and 20 µg/mL puromycin for gene knockout). The generation of the desired strains were confirmed by molecular and fluorescence-based analyses as described in the Results section.

### Scanning electron microscopy (SEM)

1.5 × 10^7^ *L. major* promastigotes were harvested and washed twice with ice-cold PBS. Cells were fixed in 2.5% glutaraldehyde at 4 °C for 1 h in the dark, followed by two washes with PBS and Milli-Q water. Cells were next resuspended in 1% osmium tetroxide solution for 20 min at room temperature. Fixed cells were washed twice with PBS and dehydrated through an ethanol gradient series (30%, 50%, 60%, and 80%), followed by resuspension in absolute ethanol. The dehydrated cell suspension was spread onto silicon wafer pieces and air-dried. Samples were further dried under vacuum in a desiccator until imaging. Prior to imaging, samples were gold-coated and examined using a scanning electron microscope (Zeiss Supra 55VP). Morphometric analysis was performed using Fiji (ImageJ).

### Analysis of *L. major* cell growth

Growth kinetics of *L. major* wild-type and LmME2^-/-^ parasites were assessed by monitoring cell proliferation over time. Promastigotes were seeded at an equal density of 5 × 10^5^ cells/mL and cultured under standard conditions. Cell growth was measured at 24, 48, and 72 h. Growth curves were generated based on cell density at each time point. To evaluate the effect of pentose phosphate pathway inhibition, parasites were treated with the 4 µM glucose-6-phosphate dehydrogenase inhibitor G6PDi, and cell growth was assessed at 72 h post-treatment. For all experiments, cell counts were determined by hemocytometer-based counting using light microscope.

### Quantification of NADP(H) and NAD(H) levels

Intracellular NADP^+^ and NADPH levels in *L. major* promastigotes were quantified using a NADP^+^/NADPH assay kit (Abcam; Cat. No. ab65349). The measurements were done according to the manufacturer’s protocol with minor modifications. Briefly, 4 × 10□ cells were harvested and washed twice with PBS, followed by resuspension in 500 µL extraction buffer. Cells were lysed by two freeze-thaw cycles with intermittent vortexing. Cell lysates were centrifuged at 14,000 × g for 10 min for remaining debris to precipitate. The clear supernatant was collected for analysis of total NADP (NADP^+^ + NADPH). For selective measurement of NADPH, an aliquot (200 µL) of the lysate was incubated at 60 °C for 30 min to decompose NADP^+^ respectively. Both heat-treated and untreated samples were loaded into a 96-well plate and incubated with cycling enzyme mix and developer solution for 3 h at room temperature. Absorbance was measured at 450 nm using a Cytation 5 multi-mode plate reader (BioTek). NADPH levels were determined directly from heat-treated samples, while NADP^+^ levels were calculated by subtracting NADPH or from total NADP. The NADPH/NADP^+^ ratio was subsequently derived. The intracellular NAD^+^ and NADH levels were measured similarly using a NAD^+^/NADH assay kit was used (Elabscience; Cat no. E-BC-K804-M).

### Measurement of intracellular reactive oxygen species (ROS)

Intracellular ROS levels in *L. major* promastigotes were measured using CM-HcDCFDA, a cell-permeable fluorogenic probe that emits fluorescence upon oxidation. Wild-type and mutant parasites were cultured in the absence or presence of 500 µM hydrogen peroxide (H_2_O_2_) for 30 min. Following treatment, cells were harvested by centrifugation to remove H2O_2_-containing medium and incubated with 10 µM CM-HcDCFDA in PBS for 15 min at 26 °C in the dark with gentle shaking. Fluorescence intensity was measured by flow cytometry using a FACSCalibur instrument (BD Biosciences). At least 10,000 events were acquired per sample. Data were analyzed using CellQuest Pro software (CellQuest Pro), and mean fluorescence intensity (FITC channel) was used as a measure of intracellular ROS levels.

### *L. major* infection of J774A.1 macrophages for parasite burden determination

J774A.1 macrophages were infected with *L. major* following previous protocols as described above. Briefly, J774A.1 macrophages were activated with 100 ng/ml liopolysaccharide for 6 h followed by addition of late stationary phase *L. major* promastigotes from 4-5 days old culture at MOI 1:30. 12 h post infection, the uninternalized parasite are washed off with PBS and fresh DMEM medium was added to the cells and further incubated for 18-24 h as per experimental requirement. The cells were washed with PBS and fixed with 1:1 (v/v) acetone:methanol and mounted in DAPI-containing mounting medium (VectaShield from Vector Laboratories) to visualise the macrophage and parasite nuclei. At least 100 macrophage nuclei were imaged from different fields using Olympus IX81 epifluorescence microscope. Parasite burden was quantified by counting the number of *L. major* nuclei around macrophage nuclei, and represented as amastigotes per 100 macrophages, plotted from three independent experiments.

### *L. major* infection of BALB/c mice

*Leishmania* infection experiments in BALB/c mice were performed as previously described (44, 67), with minor modification, at IISER Kolkata Small Animal Facility following IAEC-approved protocols. Briefly, 6-8-weeks old female BALB/c mice were infected with 5 × 10^6^ late stationary phase *L. major* promastigotes through subcutaneous injection in their left hind footpads. For uninfected control set, mice were injected similarly with PBS. Lesion development in each mouse was assessed weekly by measurement of the width and thickness of both the uninfected and infected foots of each mouse using a digital vernier caliper till 12 weeks post infection. Lesion score was calculated using the formula: [(width of infected footpad - width of uninfected footpad) × (thickness of infected footpad - thickness of uninfected footpad)]. Parasite burden in the infected mice footpads was determined by the serial dilution method (67, 68). Briefly, hind limb footpads were sequentially disinfected in HiShield PVP solution (HiMedia) and washed with distilled water followed by excision. The excised tissues were weighed and homogenised in complete M199 medium. Each tissue homogenate was serially diluted in complete M199 medium and seeded in the same medium in a 96-well cell culture plate. The number of viable parasites per milligram of tissue was estimated from the highest dilution at which parasite could be grown after a period of 10 days incubation at 26 °C. A total of 7-9 mice was used in two independent experimental sets.

### Statistical analysis

All statistical analyses were executed by Student’s t test or Mann-Whitney test. The data are represented as mean ± SD or mean ± SEM from minimum three independent experiments. P-values of ≤0.05 were considered statistically significant, with levels of statistical significance indicated as follows: *p ≤ 0.05, **p < 0.01, ***p < 0.001, ****p < 0.0001.

## Acknowledgements

The authors thank Dr. Sankar Maiti and Dr. Subhankar Dolai for sharing reagents and for providing valuable suggestions. Anti-LmxPGAM and anti-LdActin antibodies were a kind gift from Dr. Frédéric Bringaud (University of Bordeaux, France) and Dr. Amogh A. Sahasrabuddhe (CSIR-CDRI, Lucknow, India). Dr. Subrata Adak (CSIR-IICB, Kolkata) is acknowledged for sharing the CRISPR-Cas9 plasmids. This work was supported by Indian Council of Medical Research (ICMR) research grant No: 6/9-7(318)/2023-ECD-II and West Bengal DSTBT grant No: 398(Sanc.)/STBT-13015/16/2024-ST SEC awarded to RD. AP was supported by CSIR NET fellowship.

## Data and resource availability

All data generated and analyzed during this study are included in the article or supplementary material, further inquiries can be directed to the corresponding author.

## Supplementary information

**Sup. Fig. 1.**
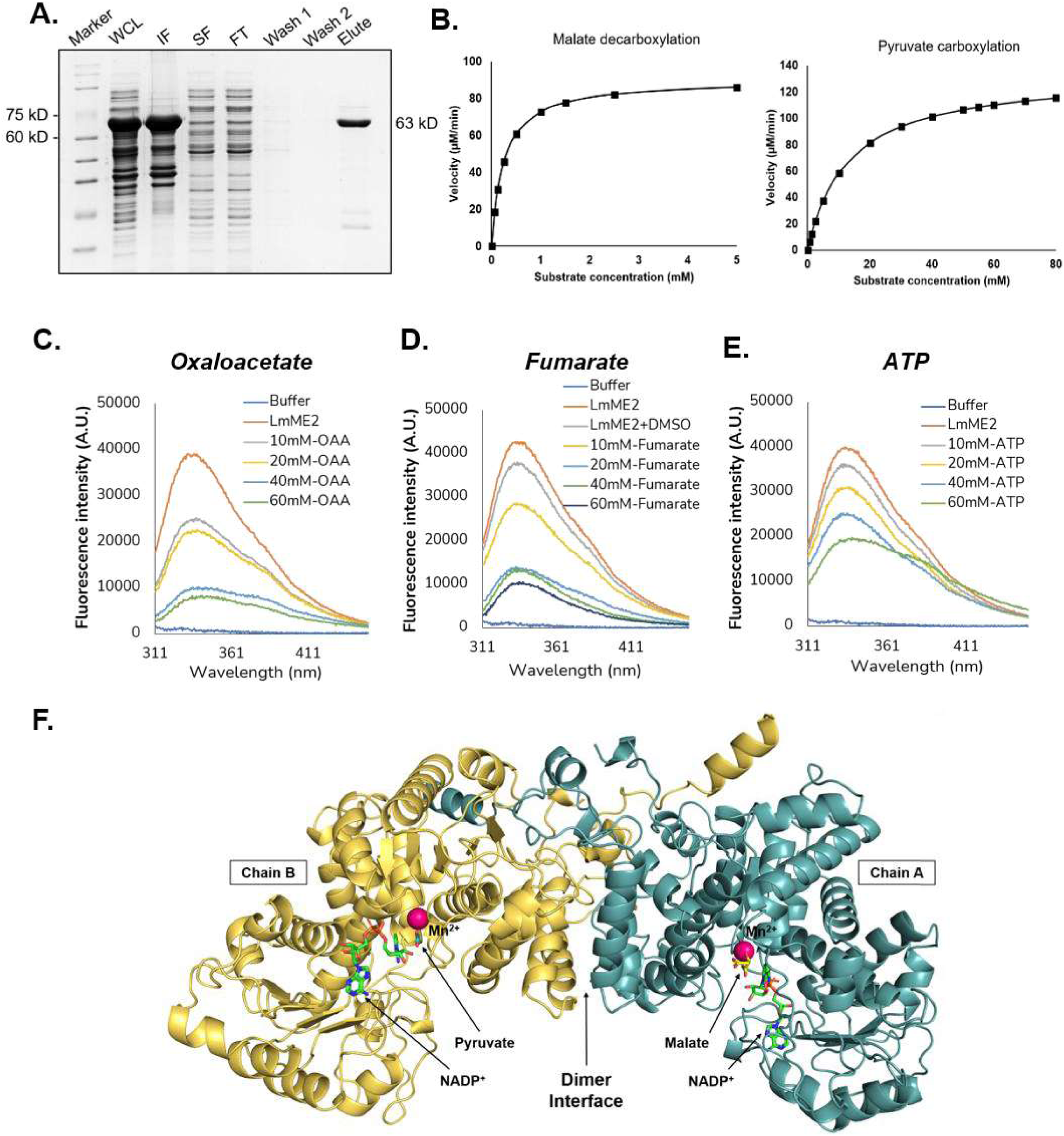
(A) Recombinant N-terminal His-tagged LmME2 (62.7 kD) was purified from *E. coli* BL21 cells by batch purification method using Ni-NTA affinity chromatography. The different fractions were loaded onto 12% SDS-PAGE gel and detected by Coomassie blue staining. The samples loaded in the lanes are as follows: WCL – whole cell lysate, IF-insoluble fraction, SF- soluble fraction, FT – flow through, Wash 1 and 2 – wash fractions (wash step was repeated twice), Elute – eluted protein. ^+^). Michaelis-Menten plots obtained for malic enzyme assay performed with 4 µg of purified LmME2 and varying concentrations of substrates malate or pyruvate and saturating concentrations of coenzymes NADP^+^ or NADPH for (B) malate decarboxylation and pyruvate carboxylation reactions respectively. Fluorescence intensity curves for tryptophan fluorescence quenching assay performed with purified recombinant LmME2 in presence of increasing concentrations of its activity regulators (C) oxaloacetate, (D) fumarate and (E) ATP. (F) AlphaFold3 prediction model for LmME2 dimer structure with the docked substrates (malate and pyruvate), the cofactor (Mn^2+^) and the coenzyme (NADP^+^).

**Sup. Fig. 2.**
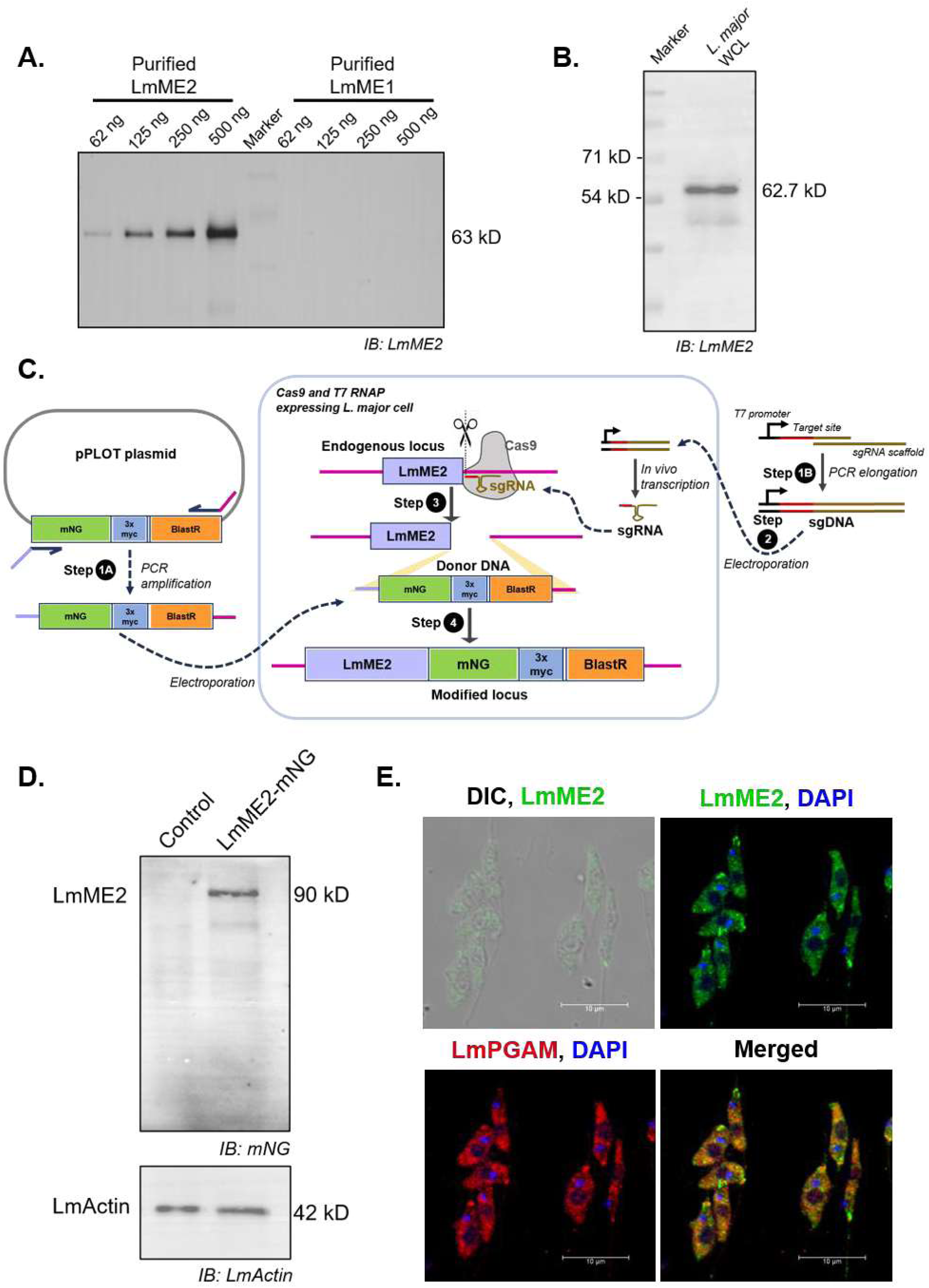
(A) Varying amounts of purified LmME2 or LmME1 (62 ng, 125 ng, 250 ng, 500 ng) or (B) 20 µg of whole cell *L. major* lysate were loaded on 12 % SDS-PAGE gel followed by western blotting with anti-LmME2 antibody to check for specificity and sensitivity of our generated polyclonal anti-sera against LmME2. (C) A schematic representation of the CRISPR-Cas9 mediated C-terminal tagging of endogenous LmME2 by mNeonGreen. Step 1: (1A) PCR amplification of donor DNA template from tagging plasmid pPLOT Blast using primers containing homologous sequences of LmME2 flanking regions (1B) PCR elongation of sgRNA template primer to form double-stranded sgRNA template DNA, capable of PAM site recognition. Step 2: Both sgRNA template DNA and donor DNA are co-electroporated into L. major promastigotes expressing SpCas9 nuclease and T7 RNA polymerase (referred to as LmCas9/T7 strain). Step 3: Cas9 directed by *in vivo* transcribed sgRNA cleaves the double strand at target site in the endogenous locus of LmME2, in its C-terminal end. Step 4: Double strand break is repaired by 30-nt flanking region containing donor DNA sequence with Blasticidin resistance antibiotic marker. Following antibiotic selection, *L. major* promastigotes with endogenously tagged LmME2 are obtained. (D) Verification of successful tagging of LmME2 (63 kD) with mNG (28 kD) by loading 15 µg of whole cell lysates of *L. major* promastigotes of strains: control (LmCas9/T7) and mNG tagged LmME2 expressing cells (LmME2-mNG) on 12% SDS-PAGE gel followed by western blotting with anti-mNG monoclonal antibody. LmActin was used as loading control. (E) Representative images of co-localization of LmME2 with known cytosolic marker *L. major* phosphoglycerate mutase (LmPGAM). Wild-type *L. major* promastigotes were co-immunostained with anti-LmME2 polyclonal antibody (green) and anti-LmPGAM polyclonal antibody (red). Nuclei were stained with DAPI (blue). The stained cells were visualised under Leica SP8 confocal microscope and the staining patterns were merged to determine colocalization. Scale bars = 10 µm.

**Sup. Fig. 3.**
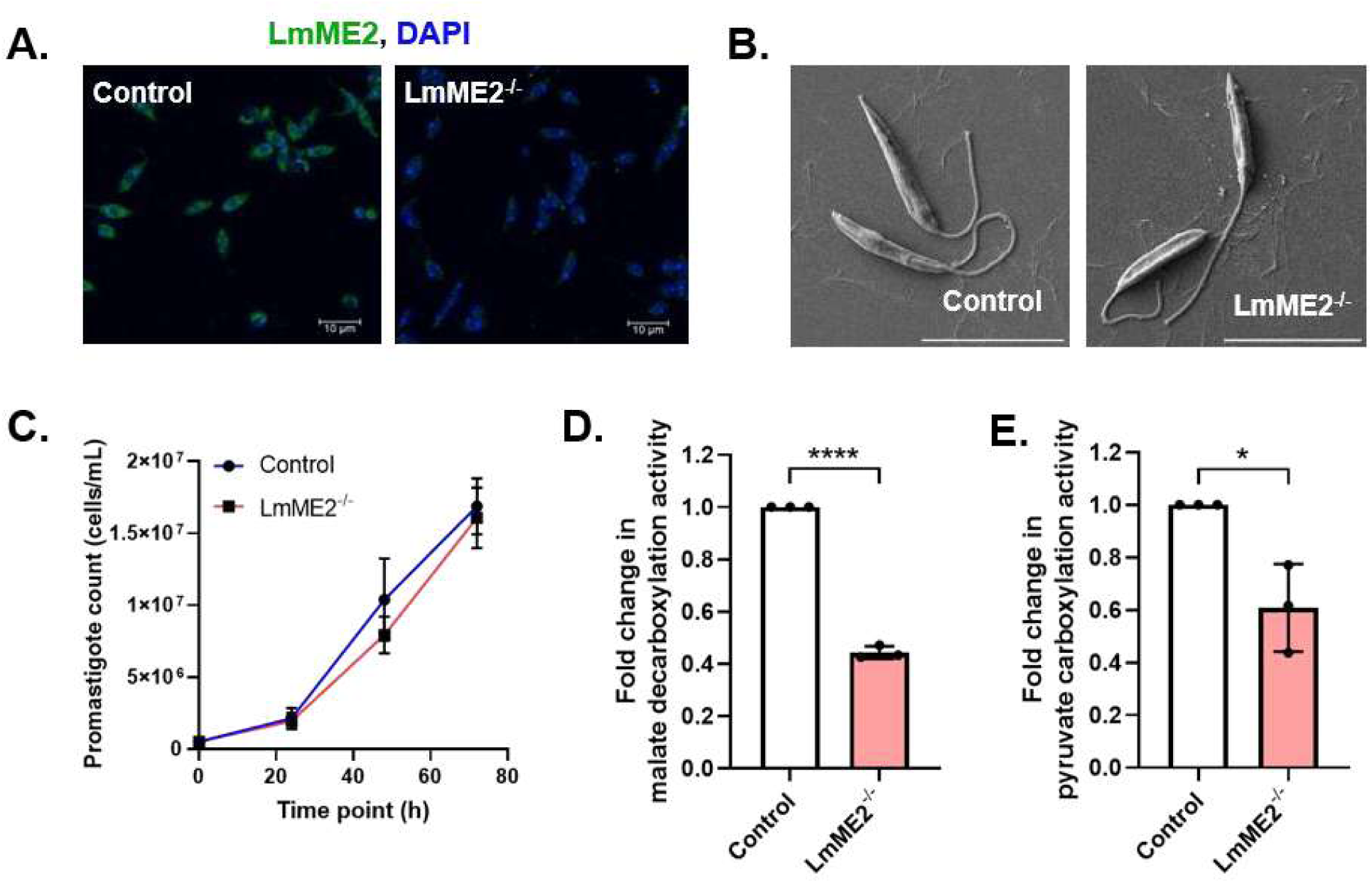
(A) Representative maximum intensity projection images of confocal microscopy of control (LmCas9/T7) and LmME2^-/-^ *L. major* promastigotes. Scale bars: 10 μm. (B) SEM images (magnification) of control (LmCas9/T7) and LmME2^-/-^ *L. major* promastigotes of late log phase. Scale bars: 10 μm. (C) Control (LmCas9/T7, blue line; round) and LmME2^-/-^ (red line; square) strains were grown for 72 h and their growth was monitored every 24 h. Each data point represents the mean±SD from 3 independent experiments. Malic enzyme assay was performed with 20 μg whole cell lysates of control and LmME2^-/-^ strains. Specific activities of both (D) malate decarboxylation and (E) pyruvate carboxylation reactions were measured. Error bars represent mean±SD of values from 3 independent experiments. Statistical significance was tested by Student’s t-test. Asterisks indicate significant difference. *P ≤ 0.05, **P ≤ 0.01 ***P ≤ 0.)01, ****P ≤ 0.0)01

**Sup. Fig. 4.**
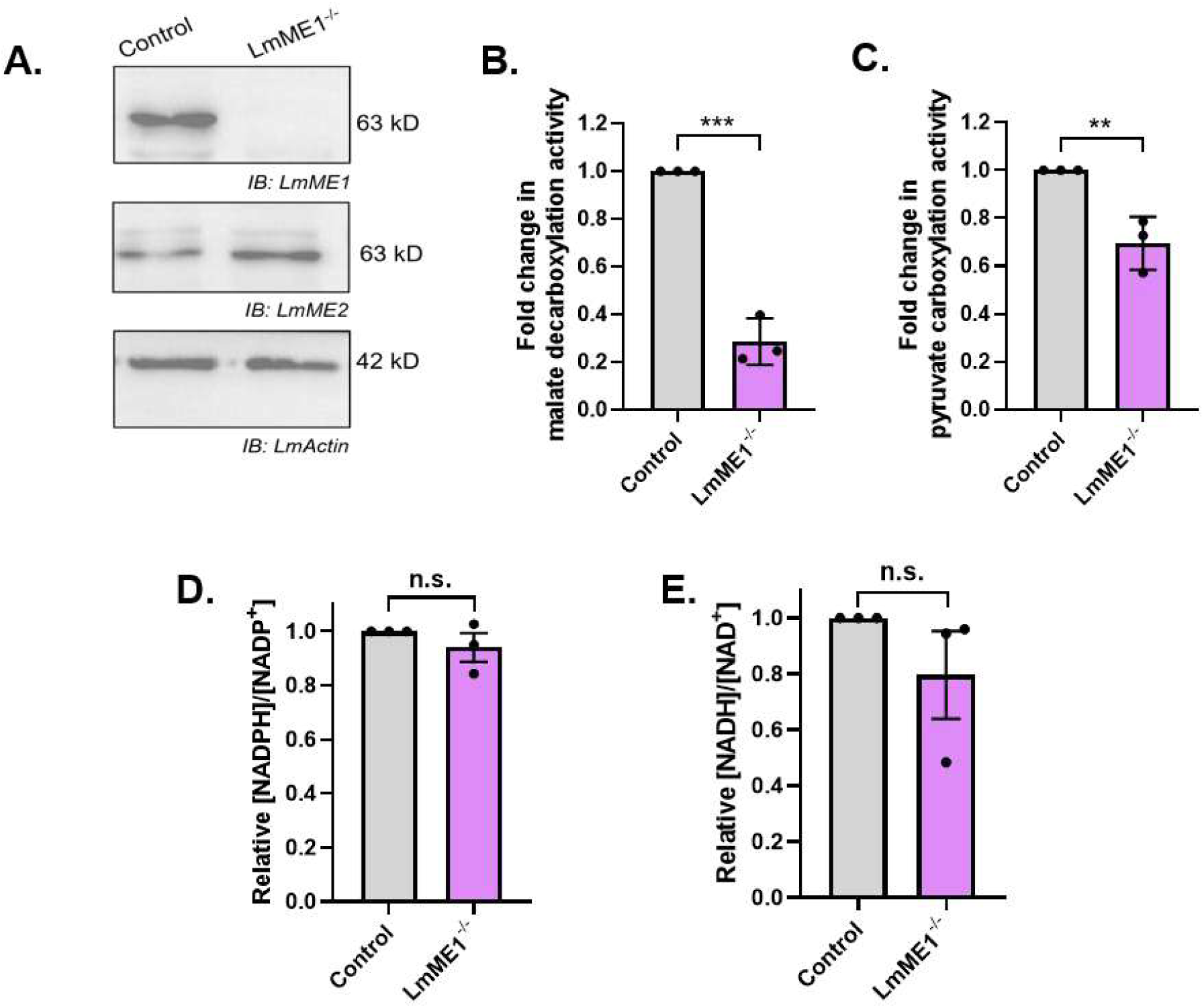
(A) Western blot images of whole cell lysates of control cells (LmCas9/T7) and LmME1^-/-^ strain with anti-LmME1 antibodies. The presence of the other isoform, LmME2 was determined by western blotting with anti-LmME2 antibody. LmActin was used as housekeeping control. Malic enzyme assay was performed with 20 μg whole cell lysates of control and LmME1^-/-^ strains to determine the specific activities of the two catalysed reaction. Bar graphs for fold change in (B) malate decarboxylation and (C) pyruvate carboxylation reactions. Error bars represent mean±SD of values from 3 independent experiments. Statistical significance was tested by unpaired Student’s t-test. NADPH, NADP^+^, NADH and NAD^+^ levels were measured for late log phase *L. major* promastigotes of control (LmCas9/T7) and LmME1^-/-^ strains. Bar graphs for relative (D) NADPH/NADP^+^ ratio, and (E) NADH/NAD^+^ ratio with respect to control (LmCas9/T7) strain. Error bars represent mean±SEM. of values from 3 independent experiments. Statistical significance was tested by Student’s t- test. Asterisks indicate significant difference. **P ≤ 0.01, n.s not significant.

**Table S1.**
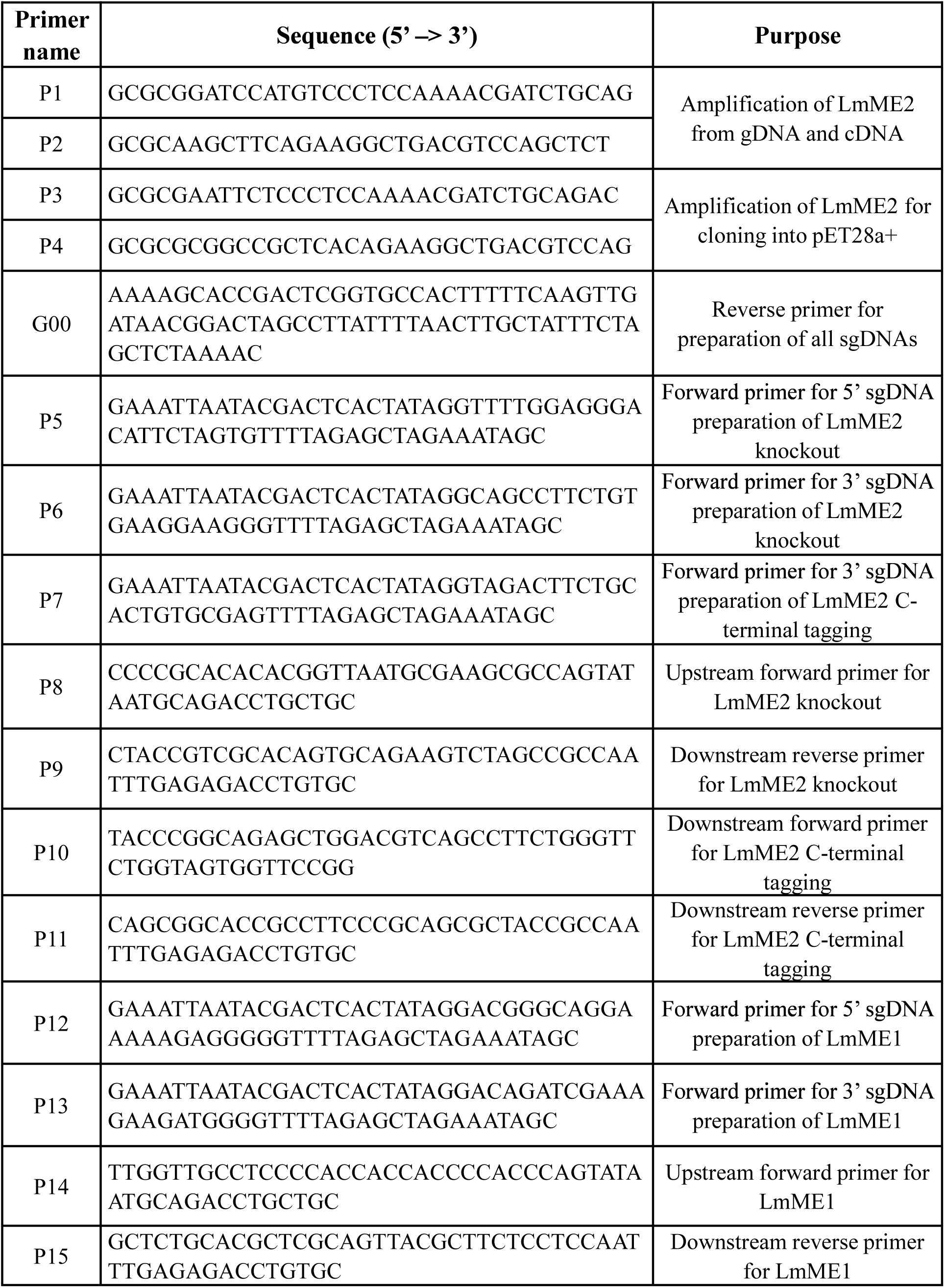
List of oligonucleotides used.

**Table S2.** List of antibodies used.

| **Antibody** | **Source** | **Catalogue No.** | **Dilution used** |
| --- | --- | --- | --- |
| Mouse anti-LmME2 | Generated in-house | - | WB - 1:4000<br>IF - 1:300 |
| Rabbit anti-mNeonGreen Tag | Cell Signaling Technology | 53061S | WB - 1:1000 |
| Rabbit anti-LmME1 | Generated in-house (21) | - | WB - 1:4000 |
| Rabbit anti-LdActin | Gifted by Dr. Amogh Sahasrabuddhe (CSIR-CDRI) | - | WB - 1:20000 |
| Rabbit anti-LmxPGAM | Gifted by Dr. Fredrich Bringaud (University of Bordeaux, France) | - | WB - 1:1000<br>IF - 1:100 |
| Goat anti-rabbit IgG Peroxidase | Sigma Aldrich | A9169 | WB - 1:4000 |
| Rabbit anti-mouse IgG Peroxidase | Sigma Aldrich | A9044 | WB - 1:4000 |
| Goat Anti-Mouse Alexa-488 | Thermo Scientific | A11001 | IF - 1:800 |
| Goat Anti-Rabbit Alexa-568 | Thermo Scientific | A11011 | IF - 1:800 |

WB- western blot, IF - Immunofluorescence

## References

1. Pareyn, M., Alves, F., Burza, S., Chakravarty, J., Alvar, J., Diro, E., Kaye, P. M., and van Griensven, J. (2025) Leishmaniasis. Nat Rev Dis Primers. 11, 81

2. Ponte-Sucre, A., Gamarro, F., Dujardin, J.-C., Barrett, M. P., López-Vélez, R., García-Hernández, R., Pountain, A. W., Mwenechanya, R., and Papadopoulou, B. (2017) Drug resistance and treatment failure in leishmaniasis: A 21st century challenge. PLOS Neglected Tropical Diseases. 11, e0006052

3. Sundar, S., and Singh, A. (2018) Chemotherapeutics of visceral leishmaniasis: present and future developments. Parasitology. 145, 481–489

4. Sacks, D., and Kamhawi, S. (2001) Molecular Aspects of Parasite-Vector and Vector-Host Interactions in Leishmaniasis. Annu. Rev. Microbiol. 55, 453–483

5. Peters, N. C., Egen, J. G., Secundino, N., Debrabant, A., Kimblin, N., Kamhawi, S., Lawyer, P., Fay, M. P., Germain, R. N., and Sacks, D. (2008) In vivo imaging reveals an essential role for neutrophils in Leishmaniasis transmitted by sand flies. Science. 321, 970–974

6. Diaz-Albiter, H., Mitford, R., Genta, F. A., Sant’Anna, M. R. V., and Dillon, R. J. (2011) Reactive Oxygen Species Scavenging by Catalase Is Important for Female Lutzomyia longipalpis Fecundity and Mortality. PLoS One. 6, e17486

7. Naderer, T., and McConville, M. J. (2008) The Leishmania–macrophage interaction: a metabolic perspective. Cellular Microbiology. 10, 301–308

8. McConville, M. J., and Naderer, T. (2011) Metabolic pathways required for the intracellular survival of Leishmania. Annu Rev Microbiol. 65, 543–561

9. Haas, A. (2007) The Phagosome: Compartment with a License to Kill. Traffic. 8, 311–330

10. Zilberstein, D., and Shapira, M. (1994) The role of pH and temperature in the development of Leishmania parasites. Annu Rev Microbiol. 48, 449–470

11. Antoine, J. C., Jouanne, C., and Ryter, A. (1990) Parasitophorous vacuoles of Leishmania amazonensis-infected macrophages maintain an acidic pH. Infect. Immun. 58, 779–787.

12. Castro, H., Tomás, A. M., and Cordeiro-da-Silva, A. (2014) Leishmania amazonensis amastigotes highly express a tryparedoxin peroxidase isoform that increases parasite resistance to macrophage antimicrobial defenses and fosters parasite virulence. PLoS Negl. Trop. Dis. 8, e3000

13. Murray, H. W., and Nathan, C. F. (1999) Macrophage microbicidal mechanisms in vivo: reactive nitrogen versus oxygen intermediates in the killing of intracellular visceral Leishmania donovani. J. Exp. Med. 189, 741–746

14. McConville, M. J., Saunders, E. C., Kloehn, J., and Dagley, M. J. (2015) Leishmania carbon metabolism in the macrophage phagolysosome- feast or famine? F1000Res. 4, 938

15. Chang, G.-G., and Tong, L. (2003) Structure and function of malic enzymes, a new class of oxidative decarboxylases. Biochemistry. 42, 12721–12733

16. Ochoa, S., Mehler, A., and Kornberg, A. (1947) REVERSIBLE OXIDATIVE DECARBOXYLATION OF MALIC ACID. Journal of Biological Chemistry. 167, 871–872

17. Murai, S., Ando, A., Ebara, S., Hirayama, M., Satomi, Y., and Hara, T. (2017) Inhibition of malic enzyme 1 disrupts cellular metabolism and leads to vulnerability in cancer cells in glucose-restricted conditions. Oncogenesis. 6, e329

18. Dey, P., Baddour, J., Muller, F., Wu, C. C., Wang, H., Liao, W. T., et al. (2017) Genomic deletion of malic enzyme 2 confers collateral lethality in pancreatic cancer. Nature. 542, 119–123.

19. Hao, G., Chen, H., Wang, L., Gu, Z., Song, Y., Zhang, H., Chen, W., and Chen, Y. Q. (2014) Role of malic enzyme during fatty acid synthesis in the oleaginous fungus Mortierella alpina. Appl Environ Microbiol. 80, 2672–2678

20. Hassel, B., and Bråthe, A. (2000) Neuronal pyruvate carboxylation supports formation of transmitter glutamate. J. Neurosci. 20, 1342–1347.

21. Mondal, D. K., Pal, D. S., Abbasi, M., and Datta, R. (2021) Functional partnership between carbonic anhydrase and malic enzyme in promoting gluconeogenesis in Leishmania major. The FEBS Journal. 288, 4129–4152

22. Liu, L., Shah, S., Fan, J., Park, J. O., Wellen, K. E., and Rabinowitz, J. D. (2016) Malic enzyme tracers reveal hypoxia-induced switch in adipocyte NADPH pathway usage. Nat Chem Biol. 12, 345–352

23. Simmen, F. A., Pabona, J. M. P., Al-Dwairi, A., Alhallak, I., Montales, M. T. E., and Simmen, R. C. M. (2023) Malic Enzyme 1 (ME1) Promotes Adiposity and Hepatic Steatosis and Induces Circulating Insulin and Leptin in Obese Female Mice. Int J Mol Sci. 24, 6613

24. Pound, K. M., Sorokina, N., Ballal, K., Berkich, D. A., Fasano, M., Lanoue, K. F., Taegtmeyer, H., O’Donnell, J. M., and Lewandowski, E. D. (2009) Substrate-enzyme competition attenuates upregulated anaplerotic flux through malic enzyme in hypertrophied rat heart and restores triacylglyceride content: attenuating upregulated anaplerosis in hypertrophy. Circ Res. 104, 805–812

25. Hsieh, J.-Y., Chen, K.-C., Wang, C.-H., Liu, G.-Y., Ye, J.-A., Chou, Y.-T., Lin, Y.-C., Lyu, C.-J., Chang, R.-Y., Liu, Y.-L., Li, Y.-H., Lee, M.-R., Ho, M.-C., and Hung, H.-C. (2023) Suppression of the human malic enzyme 2 modifies energy metabolism and inhibits cellular respiration. *Commun*. Biol. 6, 548

26. Chen, K.-C., Hsiao, I.-H., Huang, Y.-N., Chou, Y.-T., Lin, Y.-C., Hsieh, J.-Y., Chang, Y.-L., Wu, K.-H., Liu, G.-Y., and Hung, H.-C. (2023) Targeting human mitochondrial NAD(P)+-dependent malic enzyme (ME2) impairs energy metabolism and redox state and exhibits antileukemic activity in acute myeloid leukemia. Cell Oncol (Dordr). 46, 1301–1316

27. Hasan, N. M., Longacre, M. J., Stoker, S. W., Kendrick, M. A., and MacDonald, M. J. (2015) Mitochondrial malic enzyme 3 is important for insulin secretion in pancreatic β-cells. Mol Endocrinol. 29, 396–410

28. Ren, J. G., Seth, P., Clish, C. B., Lorkiewicz, P. K., Higashi, R. M., Lane, A. N., et al. (2014) Knockdown of malic enzyme 2 suppresses lung tumor growth, induces differentiation and impacts PI3K/AKT signaling. Sci. Rep. 4, 5414.

29. Jiang, P., Du, W., Mancuso, A., Wellen, K. E., and Yang, X. (2013) Reciprocal regulation of p53 and malic enzymes modulates metabolism and senescence. Nature. 493, 689–693

30. Drincovich, M. F., Casati, P., and Andreo, C. S. (2001) NADP-malic enzyme from plants: a ubiquitous enzyme involved in different metabolic pathways. FEBS Letters. 490, 1–6

31. Tronconi, M. A., Fahnenstich, H., Gerrard Weehler, M. C., Andreo, C. S., Flügge, U.-I., Drincovich, M. F., and Maurino, V. G. (2008) Arabidopsis NAD-malic enzyme functions as a homodimer and heterodimer and has a major impact on nocturnal metabolism. Plant Physiol. 146, 1540–1552

32. Saigo, M., Tronconi, M. A., Gerrard Wheeler, M. C., Alvarez, C. E., Drincovich, M. F., and Andreo, C. S. (2013) Biochemical approaches to C4 photosynthesis evolution studies: the case of malic enzymes decarboxylases. Photosynth Res. 117, 177–187

33. Bologna, F. P., Andreo, C. S., and Drincovich, M. F. (2007) Escherichia coli malic enzymes: two isoforms with substantial differences in kinetic properties, metabolic regulation, and structure. J Bacteriol. 189, 5937–5946

34. Lerondel, G., Doan, T., Zamboni, N., Sauer, U., and Aymerich, S. (2006) YtsJ has the major physiological role of the four paralogous malic enzyme isoforms in Bacillus subtilis. J Bacteriol. 188, 4727–4736

35. Boles, E., de Jong-Gubbels, P., and Pronk, J. T. (1998) Identification and characterization of MAE1, the Saccharomyces cerevisiae structural gene encoding mitochondrial malic enzyme. J Bacteriol. 180, 2875–2882

36. Zhu, B. H., Zhang, R. H., Lv, N. N., Yang, G. P., Wang, Y. S., and Pan, K. H. (2018) The role of malic enzyme on promoting total lipid and fatty acid production in Phaeodactylum tricornutum. Front. Plant Sci. 9, 826.

37. Leroux, A. E., Maugeri, D. A., Opperdoes, F. R., Cazzulo, J. J., and Nowicki, C. (2011) Comparative studies on the biochemical properties of the malic enzymes from Trypanosoma cruzi and Trypanosoma brucei. FEMS Microbiol Lett. 314, 25–33

38. Allmann, S., Morand, P., Ebikeme, C., Gales, L., Biran, M., Hubert, J., Brennand, A., Mazet, M., Franconi, J.-M., Michels, P. A. M., Portais, J.-C., Boshart, M., and Bringaud, F. (2013) Cytosolic NADPH Homeostasis in Glucose-starved Procyclic Trypanosoma brucei Relies on Malic Enzyme and the Pentose Phosphate Pathway Fed by Gluconeogenic Flux *. Journal of Biological Chemistry. 288, 18494–18505

39. Giordana, L., Sosa, M. H., Leroux, A. E., Mendoza, E. F. R., Petray, P., and Nowicki, C. (2018) Molecular and functional characterization of two malic enzymes from Leishmania parasites. Mol Biochem Parasitol. 219, 67–76

40. Camacho, E., González-de la Fuente, S., Solana, J. C., Rastrojo, A., Carrasco-Ramiro, F., Requena, J. M., and Aguado, B. (2021) Gene annotation and transcriptome delineation on a de novo genome assembly for the reference Leishmania major Friedlin strain. Genes (Basel) 12, 1359

41. Yang, Z., Lanks, C. W., and Tong, L. (2002) Molecular mechanism for the regulation of human mitochondrial NAD(P)+-dependent malic enzyme by ATP and fumarate. Structure. 10, 951–960

42. McKenna, M. C., Tildon, J. T., Stevenson, J. H., Huang, X., and Kingwell, K. G. (1995) Regulation of mitochondrial and cytosolic malic enzymes from cultured rat brain astrocytes. Neurochem Res. 20, 1491–1501

43. Guerra, D. G., Vertommen, D., Fothergill-Gilmore, L. A., Opperdoes, F. R., and Michels, P. A. M. (2004) Characterization of the cofactor-independent phosphoglycerate mutase from Leishmania mexicana mexicana. Histidines that coordinate the two metal ions in the active site show different susceptibilities to irreversible chemical modification. Eur J Biochem. 271, 1798–1810

44. Samanta, S., Banerjee, S., and Datta, R. (2025) A secreted *Leishmania* metalloprotease manipulates host iron regulation by targeting the DICER1–miRNA pathway. Journal of Biological Chemistry. 301, 110851

45. Seth, A., Das, A., and Datta, R. (2025) Identification of a basal body-localized epsilon-tubulin in Leishmania. FEBS Letters. 599, 1569–1581

46. Szöőr, B., Simon, D. V., Rojas, F., Young, J., Robinson, D. R., Krüger, T., Engstler, M., and Matthews, K. R. (2019) Positional Dynamics and Glycosomal Recruitment of Developmental Regulators during Trypanosome Differentiation. mBio. 10, e00875–19

47. Oka, S.-I., Titus, A. S., Zablocki, D., and Sadoshima, J. (2023) Molecular properties and regulation of NAD+ kinase (NADK). Redox Biol. 59, 102561

48. Chen, X., Zhang, Y., Song, D., Gui, F., Cao, Y., Hong, Y., Chen, R., Song, Y., Di, C., Yang, J., and Tan, X. (2025) NADK Governs Ferroptosis Susceptibility by Orchestrating NADPH Homeostasis. Antioxidants (Basel). 14, 1396

49. Maugeri, D. A., and Cazzulo, J. J. (2004) The pentose phosphate pathway in Trypanosoma cruzi. FEMS Microbiol Lett. 234, 117–123

50. Allmann, S., Morand, P., Ebikeme, C., Gales, L., Biran, M., Hubert, J., Brennand, A., Mazet, M., Franconi, J.-M., Michels, P. A. M., Portais, J.-C., Boshart, M., and Bringaud, F. (2013) Cytosolic NADPH homeostasis in glucose-starved procyclic Trypanosoma brucei relies on malic enzyme and the pentose phosphate pathway fed by gluconeogenic flux. J Biol Chem. 288, 18494–18505

51. Ghergurovich, J. M., García-Cañaveras, J. C., Wang, J., Schmidt, E., Zhang, Z., TeSlaa, T., Patel, H., Chen, L., Britt, E. C., Piqueras-Nebot, M., Gomez-Cabrera, M. C., Lahoz, A., Fan, J., Beier, U. H., Kim, H., and Rabinowitz, J. D. (2020) A small molecule G6PD inhibitor reveals immune dependence on pentose phosphate pathway. Nat Chem Biol. 16, 731–739

52. Watermann, P., Arend, C., and Dringen, R. (2023) G6PDi-1 is a Potent Inhibitor of G6PDH and of Pentose Phosphate pathway-dependent Metabolic Processes in Cultured Primary Astrocytes. Neurochem Res. 48, 3177–3189

53. Ivens, A. C., Peacock, C. S., Worthey, E. A., Murphy, L., Aggarwal, G., Berriman, M., Sisk, E., Rajandream, M.-A., Adlem, E., Aert, R., Anupama, A., Apostolou, Z., Attipoe, P., Bason, N., Bauser, C., Beck, A., Beverley, S. M., Bianchettin, G., Borzym, K., Bothe, G., Bruschi, C. V., Collins, M., Cadag, E., Ciarloni, L., Clayton, C., Coulson, R. M. R., Cronin, A., Cruz, A. K., Davies, R. M., De Gaudenzi, J., Dobson, D. E., Duesterhoeft, A., Fazelina, G., Fosker, N., Frasch, A. C., Fraser, A., Fuchs, M., Gabel, C., Goble, A., Goffeau, A., Harris, D., Hertz-Fowler, C., Hilbert, H., Horn, D., Huang, Y., Klages, S., Knights, A., Kube, M., Larke, N., Litvin, L., Lord, A., Louie, T., Marra, M., Masuy, D., Matthews, K., Michaeli, S., Mottram, J. C., Müller-Auer, S., Munden, H., Nelson, S., Norbertczak, H., Oliver, K., O’neil, S., Pentony, M., Pohl, T. M., Price, C., Purnelle, B., Quail, M. A., Rabbinowitsch, E., Reinhardt, R., Rieger, M., Rinta, J., Robben, J., Robertson, L., Ruiz, J. C., Rutter, S., Saunders, D., Schäfer, M., Schein, J., Schwartz, D. C., Seeger, K., Seyler, A., Sharp, S., Shin, H., Sivam, D., Squares, R., Squares, S., Tosato, V., Vogt, C., Volckaert, G., Wambutt, R., Warren, T., Wedler, H., Woodward, J., Zhou, S., Zimmermann, W., Smith, D. F., Blackwell, J. M., Stuart, K. D., Barrell, B., and Myler, P. J. (2005) The genome of the kinetoplastid parasite, Leishmania major. Science. 309, 436–442

54. Bringaud, F., Müller, M., Cerqueira, G. C., Smith, M., Rochette, A., El-Sayed, N. M. A., Papadopoulou, B., and Ghedin, E. (2007) Members of a large retroposon family are determinants of post-transcriptional gene expression in Leishmania. PLoS Pathog. 3, 1291–1307

55. Alonso, G., Rastrojo, A., López-Pérez, S., Requena, J. M., and Aguado, B. (2016) Resequencing and assembly of seven complex loci to improve the Leishmania major (Friedlin strain) reference genome. Parasit Vectors. 9, 74

56. Zhu, Y., Gu, L., Lin, X., Liu, C., Lu, B., Cui, K., Zhou, F., Zhao, Q., Prochownik, E. V., Fan, C., and Li, Y. (2020) Dynamic Regulation of ME1 Phosphorylation and Acetylation Affects Lipid Metabolism and Colorectal Tumorigenesis. Molecular Cell. 77, 138–149.e5

57. Teng, P., Cui, K., Yao, S., Fei, B., Ling, F., Li, C., and Huang, Z. (2024) SIRT5- mediated ME2 desuccinylation promotes cancer growth by enhancing mitochondrial respiration. Cell Death Differ. 31, 65–77

58. Li, C., Ge, C., Wang, Q., Teng, P., Jia, H., Yao, S., and Huang, Z. (2025) Sirtuin 3-mediated delactylation of malic enzyme 2 disrupts redox balance and inhibits colorectal cancer growth. Cell Oncol. 48, 979–990

59. Harding, C. J., Cadby, I. T., Moynihan, P. J., and Lovering, A. L. (2021) A rotary mechanism for allostery in bacterial hybrid malic enzymes. Nat Commun. 12, 1228

60. Su, K.-L., Chang, K.-Y., and Hung, H.-C. (2009) Effects of structural analogues of the substrate and allosteric regulator of the human mitochondrial NAD(P)+-dependent malic enzyme. Bioorg Med Chem. 17, 5414–5419

61. Xie, N., Zhang, L., Gao, W., Huang, C., Huber, P. E., Zhou, X., Li, C., Shen, G., and Zou, B. (2020) NAD+ metabolism: pathophysiologic mechanisms and therapeutic potential. Signal Transduct Target Ther. 5, 227

62. Pal, D. S., Mondal, D. K., and Datta, R. (2015) Identification of Metal Dithiocarbamates as a Novel Class of Antileishmanial Agents. Antimicrob Agents Chemother. 59, 2144–2152

63. Żelewski, M., and Świerczyński, J. (1991) Malic enzyme in human liver. European Journal of Biochemistry. 201, 339–345

64. Lowry, O. H., Rosebrough, N. J., Farr, A. L., and Randall, R. J. (1951) Protein measurement with the Folin phenol reagent. J Biol Chem. 193, 265–275

65. Pilar, A. V. C., Madrid, K. P., and Jardim, A. (2008) Interaction of *Leishmania* PTS2 receptor peroxin 7 with the glycosomal protein import machinery. Molecular and Biochemical Parasitology. 158, 72–81

66. Beneke, T., Madden, R., Makin, L., Valli, J., Sunter, J., and Gluenz, E. (2017) A CRISPR Cas9 high-throughput genome editing toolkit for kinetoplastids. R Soc Open Sci. 4, 170095

67. Banerjee, S., and Datta, R. (2023) Localized *Leishmania major* infection disrupts systemic iron homeostasis that can be controlled by oral iron supplementation. Journal of Biological Chemistry. 299, 105064

68. Sacks, D. L., and Melby, P. C. (1998) Animal Models for the Analysis of Immune Responses to Leishmaniasis. Current Protocols in Immunology. 28, 19.2.1-19.2.20

